# Modular structure of human olfactory receptor codes reflects the bases of odor perception

**DOI:** 10.1101/525287

**Authors:** Ji Hyun Bak, Seogjoo J. Jang, Changbong Hyeon

## Abstract

The circuits of olfactory signaling are reminiscent of complex computational devices. The olfactory receptor code, which represents the responses of receptors elicited by olfactory stimuli, is effectively an input code for the neural computation of odor sensing. Here, analyzing a recent dataset of the odorant-dependent responses of human olfactory receptors (ORs), we show that the space of human olfactory receptor codes is partitioned into a modular structure where groups of receptors are “labeled” for key olfactory features. Our analysis reveals a low-dimensional structure in the space of human odor perception, with the receptor groups as the bases to represent major features in the perceptual odor space. These findings provide a novel evidence that some fundamental olfactory features are already hard-coded at the level of ORs, separately from the higher-level neural circuits.

## INTRODUCTION

Olfaction is a sensory process that captures environmental signals by detecting molecular stimuli through a repertoire of olfactory receptors (ORs). Although the full processing of olfactory information is realized in the complicated neural circuits [1], the first step of olfactory sensing involves a selective binding of odorants to the cognate ORs, which is biochemical in nature [2]. The binding elicits an array of downstream responses in the corresponding olfactory receptor neurons (ORNs) [3]. The response pattern encoded into ORs or ORNs, termed the olfactory *receptor code* [4], provides the first neural representation of an odor and is essential in the early stage of olfactory sensing.

Lying at a crucial juncture in the flow of olfactory information, the receptor code bridges between the molecular and neural spaces. An olfactory stimulus in the *molecular space*, representing the physiochemical properties of the odorants, is translated into the *receptor code space* to form an “input code”. The information is then processed through the higher-order *neural spaces*, and eventually evokes the sense of smell that constitutes the *perceptual odor space* (see Fig. 1). Recent insights into the processing of olfactory information were gained mostly based on the olfactory sensing of non-human species [5–7] or through theoretical studies [8, 9]. A separate branch of research attempts to relate the molecular space directly with the perceptual odor space [10, 11]. However, the goal of understanding the principles of olfactory computation in humans faces a fundamental difficulty due to our limited access to the neural circuitry in the human brain.

**Figure 1:**
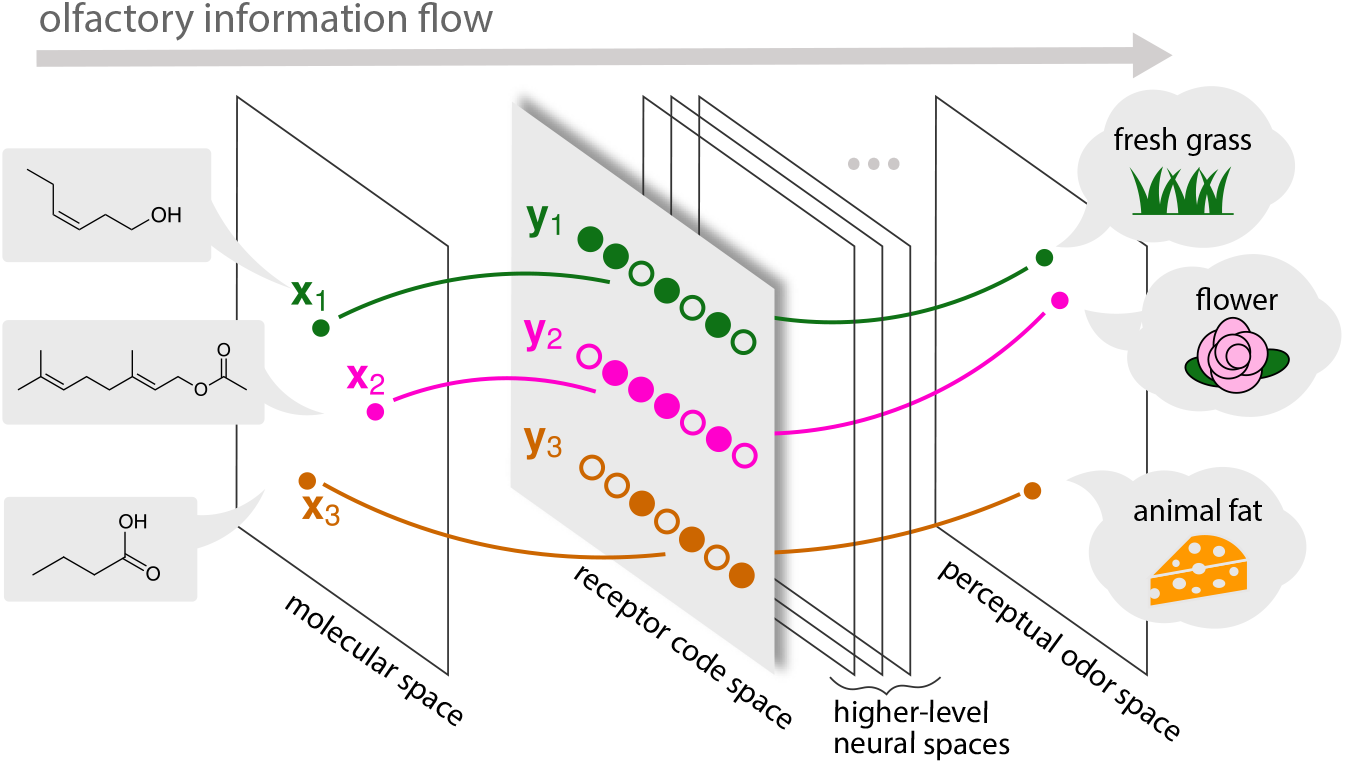
Schematic of olfactory information flow. The *molecular space*, representing physiochemical features of odorant molecules, is first recognized by OR repertoire and encoded into the *receptor code space*. The information stored in the receptor codes is modified while it goes through the higher-level *neural spaces* and finally projected on the *perceptual odor space*, and sensed as a smell.

Here, we show that a critical progress can still be made by a systematic analysis of the receptor code space. Whereas the OR repertoire is known to encode the input signal from odorants like a piano keyboard that combinatorially produces a variety of musical chords [4], it may be further “formatted” to facilitate the processing of relevant information, perhaps by appropriately reflecting the structure of the perceptual odor space. Specifically, we analyze a set of receptor codes for different odorants, extracted from the biochemical measurement of downstream responses of the human ORs [12–14]. Because the receptor code is implemented by the many-to-many pairwise interaction between odorants and ORs [4, 15], we treat the interaction of each odorant-receptor pair as either “on” (responding) or “off” (not responding), and analyze the binary pattern of pairwise interaction, considering it a zeroth-order representation of the receptor code. Our analysis of overlap between distinct receptor codes (receptor code redundancy) reveals the intrinsic structure of the human receptor code space, which is also correlated with the set of coarse-grained features in the perceptual odor space. This suggests that the processing of olfactory information is already at work in the receptor space.

## RESULTS

We present a quantitative analysis of the pattern of odorant-receptor interactions, employing a dataset that reports the responses of 303 receptors against 89 odorants [14]. The state of all *N* = 303 receptors can be represented in terms of an *N*-dimensional binary vector **y**, where *y_i_* = 1 if the *i*-th receptor is “on”, and *y_i_* = 0 if it is “off”. For a given odor **x**, the corresponding *receptor code* **y** represents the odor **x** in the *N*-dimensional binary receptor code space.

We collected all 535 pairwise interactions reported in the dataset, including 60 de-activations [16], and visualized the receptor codes in the form of an interaction network (Fig. 2a). In this interaction network, each node is either an odorant or a receptor, and each edge connects an interacting odorant-receptor pair. For a more detailed version of the interaction network with varying parameters, see Fig. S1.

**Figure 2:**
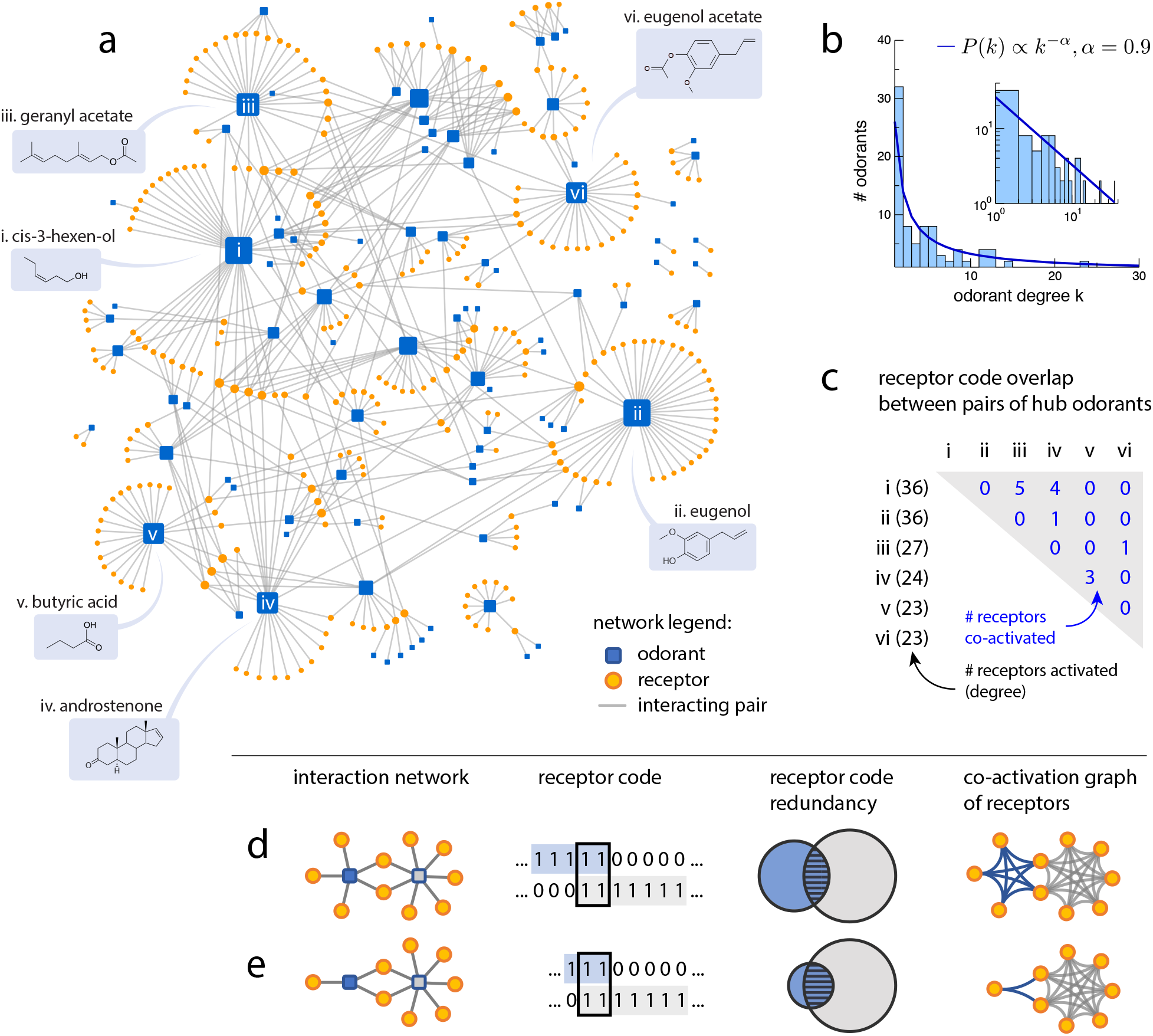
Odorant-receptor interaction network and receptor codes. **(a)** The odorant-receptor interaction network that visualizes all the interactions between 89 odorants (blue squares) and 303 receptors (orange circles) from the dataset [14]. An edge is drawn when odorant and receptor is interacting. Node size reflects the degree; six highest-degree odorants are marked. Also see Fig. S1 for an extended display with all odorant labels, and with edge attributes for pairwise interaction properties. **(b)** The histogram of odorant degrees. The degree distribution is fitted to *P*(*k*) ~ *k*^−*α*^ with *α* = 0.9 (*R*^2^ = 0.74 from a linear regression of log *P*(*k*)vs. log *k*), and has an average value 〈*k*〉 ≃ 6 (also see Fig. S2). **(c)** The overlaps between the receptors codes for the six hub odorants identified in **a** (also see Fig. S3). **(d-e)** Illustration of the idea of receptor code redundancy. The first two columns show two equivalent representations of the receptor code space, as an interaction network and as a set of binary vectors. We consider the receptor code redundancy of a target odorant (blue) with respect to a background odorant (gray). In the third column, the receptor code redundancy is illustrated with Venn diagrams of interacting receptors. The redundancy of adding the target odorant in the presence of the background odorant is represented by the relative size of overlap (striped area) with respect to the total size of target signal (blue-shaded area). Although the number of overlapping receptors is the same in both cases, the normalized redundancy is smaller in **d** (two out of five) than in **e** (two out of three). In the last column, the interaction network is projected to a co-activation graph of receptors.

### Interaction network reveals odorant hubs with non-redundant receptor codes

In the odorant-OR interaction network (Fig. 2a), the number of edges attached to an odorant node indicates the number of receptors that recognize this odorant. We call this number the *degree* of the odorant node, and denote it by *k*_λ_, where λ is the index for the odorant (also see Methods).

We observe two properties from the statistics of odorant degrees. First, the receptor-space representation of single odorants is sparse: On average, an odorant is recognized by only 6 (〈*k*〉) ≃ 6) out of *N* = 303 receptors, which amounts to 2% of the receptor space. The sparsity observed in the network is consistent with previous reports from the ORN responses [17, 18]. Second, the degrees of the odorants in the dataset are non-uniformly distributed with a heavy tail, which can be approximately fitted to *P*(*k*) ~ *k*^−*α*^ with *α* ≈ 0.9 (Fig. 2b). Such distribution allows us to identify a small number of high-degree odorant “hubs”. The six highest-degree hub odorants are labeled in Fig. 2a. It is worth noting that the receptor codes for the hub odorants are highly non-redundant with one another (Fig. 2c); among the six highest-degree odorants, 10 out of all 15 pairs have completely disjoint sets of receptor codes. In particular, despite significant similarity in chemical structures, there is no co-activated OR between eugenol and eugenol acetate, which differ in a single functional group (hydroxy group versus acetate in the aromatic ring). The overlap of receptor codes among the hub odorants is significantly smaller than what is expected from a random mixture of odorants whose receptor codes cover the same number of receptors (*p* < 0.02; also see Methods and Fig. S3).

We introduce an idea of *receptor code redundancy*, a quantitative measure of (dis-)similarity between the olfactory responses of two distinct odorants. If two odors have exactly the same receptor codes, they are perceived as the same olfactory signal. We hypothesize that two odors are poorly distinguishable when their receptor codes have a higher redundancy. For example, consider a discriminatory task where the goal is to detect a target odorant in the presence of a constant background odor [19, 20]. When there was no background odor, the signal would elicit responses in |**y**_targ_| receptors, where **y**_targ_ is the receptor code for the target odorant. But when some receptors also respond to the background, **y**_bg_, the states of the overlapping receptors would not change upon the addition of the target odorant; the receptor code overlap |**y**_targ_ ∩ **y**_bg_ | reduces the effective size of the response (also see Fig. 2d-e). Therefore we define the receptor code redundancy in terms of the fraction χ = |**y**_targ_ ∩ **y**_bg_ | / |**y**_targ_|. If this fraction is small, as in Fig. 2d, the target odorant is deemed unambiguously detected even in the presence of the background odorant.

Taken together, the single-odorant representation in the receptor space is sparse and non-uniform, giving rise to high-degree “hub” odorants. The receptor code is almost non-redundant among these hub odorants, whereas the odorants that have a greater receptor code redundancy with a hub odorant are grouped around it. As presented below, analysis of the receptor code redundancy enables us to decipher the structure of the receptor code space more quantitatively.

### The receptor code space is naturally partitioned to show a modular structure

The grouped structure is better manifested when the interaction network is projected to the space of receptors. Here we consider the *co-activation graph* of receptors, which inherits all receptor nodes in the original interaction network; two receptor nodes are connected by an edge if they share a common odorant in the original network (Fig. 2d-e, last column). Of particular interest in the coactivation graph are the “receptor *cliques”*, or the groups of receptors that are co-activated by the same odorants. Because each pair of receptors interacting with a shared odorant λ is connected in the co-activation graph of receptors, the set of all receptors that interact with a given odorant always form a clique. Given the particular structure of the interaction network, with hub odorants with largely non-redundant receptor codes (Fig. 3a), the co-activation graph of receptors is bound to have large and mostly non-overlapping cliques that are associated to the hub odorants (Fig. 3b). We use the receptor cliques to *partition* the receptor space into non-overlapping groups, such that each receptor group is associated to a shared odorant; when a receptor is a part of more than one receptor clique, it is assigned to the larger clique (Fig. 3c).

**Figure 3:**
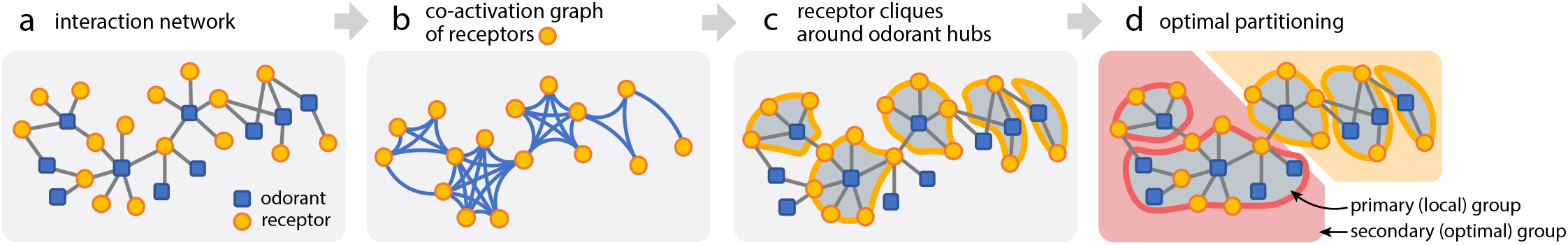
Schematics for the partitioning of receptor code. (**a**) Given an interaction network with particular statistics of the hub odorants, (**b**) the modular structure is revealed more clearly in the co-activation pattern of receptors. (**c**) We characterize the local structure from the receptor groups around the odorant hubs, and (**d**) merge the local groups to obtain the best partitioning of the receptor code. Also see Fig. S4 for a more detailed illustration.

The receptor groups can be used to group odorants based on their receptor codes. The idea is to construct odorant groups such that an odorant λ belongs to an odorant group Λ if its receptor code redundancy with respect to the group Λ, χ_λ|Λ_ = |**y**_λ_ ∩ **y**_Λ_| / |**y**_λ_|, is large (see Methods). First, we used the receptor groups to determine the reference receptor codes for odorant groups (**y**_Λ_’s). Specifically, for each receptor group Λ (note that the group index is shared between the receptors and the odorants), we make a binary vector **y**_Λ_ such that [**y**_Λ_]_*i*_ = 1 if and only if receptor *i* belongs to the group Λ. Then we assigned each odorant to the group Λ where the receptor code redundancy χ_λ|Λ_ is maximized. This results in a simultaneous partitioning of odorants and receptors (Fig. 3d, local groups).

The local groups emerge naturally from the particular statistics of the receptor codes with hub odorants, and capture the local coactivation pattern; receptors are grouped together if they are coactivated by the same odorant. If we were to characterize the overall structure of the receptor code space in terms of the groups, however, the pairs that are *not* grouped together also carry important information. In our case, a grouping is “good” if receptors in the same group respond to the same sets of odorants, *and* receptors in different groups to different odorants. Now we step further to obtain an optimal grouping of the receptor code space based on the co-activation patterns: we carry out a *secondary* grouping of the receptors by merging the *primary* groups obtained above by identifying receptor cliques. We merge a pair of primary groups when their overlap in the co-activation matrix is larger than a threshold (see Methods for details), where the threshold for merging is determined to maximize the goodness of grouping (Fig. S5). Because we partition the odorants and the receptors simultaneously, a merging of receptor groups automatically leads to a merging of the corresponding odorant groups (Fig. 3d).

Application of the grouping procedure to the human olfactory receptor codes results in the sorted interaction matrix in Fig. 4, where each row represents the receptor code for a given odorant. The rows and columns of the interaction matrix are sorted according to the orders in the respective odorant/receptor groups (Fig. 4a-b). The interaction matrix has a roughly block-diagonal form (Fig. 4c), with much less significant contributions from the off-diagonal elements. Notice that we have already arranged the interaction network layout to represent a grouped structure, which is visualized more clearly in Fig. 5, where colored territories (Groups 1–6) represent the best partitioning of the receptor code space. The feasibility of grouping reflects a special structure of the receptor code; such clear grouping is not obtained from a random network with the same degree statistics. Specifically, we randomize the interaction matrix while preserving (i) the number of interactions, (ii) the number of interactions for each odorant (degree) and (iii) the primary grouping structure. In all three cases, it was highly unlikely to find a grouping as good as or better than what we observed from the human receptor code data (*p* ≤ 0.002; see Methods and Fig. S6).

**Figure 4:**
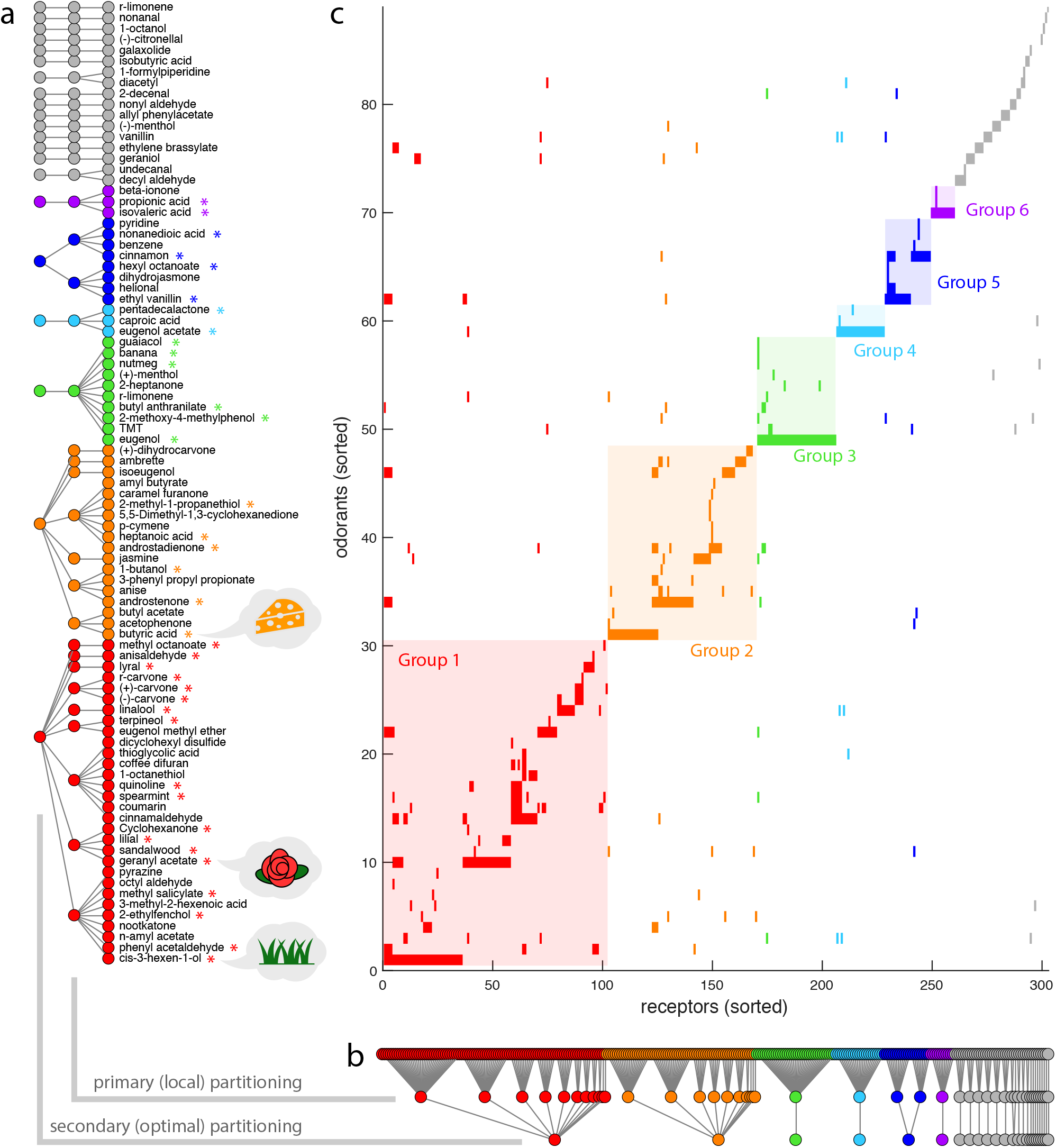
Simultaneous partitioning of odorant and receptors, based on the co-activation pattern and the receptor code redundancy. The primary and secondary groups are shown for (**a**) the odorants and (**b**) the receptors, where the indices are simultaneously sorted by the group rank. Also see Fig.S5 and Fig.S6 for validations. (**c**) The rows and columns of interaction matrix is consequently sorted by the odorant and receptor indices. Each row represents the receptor code for the corresponding odorant. The six largest secondary groups are colored across all panels. Diagonal-block shades in the interaction matrix are added for better visualization of the simultaneous partitioning of odorants and receptors. In (**a**), odorants with odor qualities consistent with the characteristic olfactory feature of the corresponding secondary group (plant-like for Group 1; animal-like for Groups 2 and 6; culinary for Groups 3, 4 and 5) are marked with asterisks. Also see Table S1.

**Figure 5:**
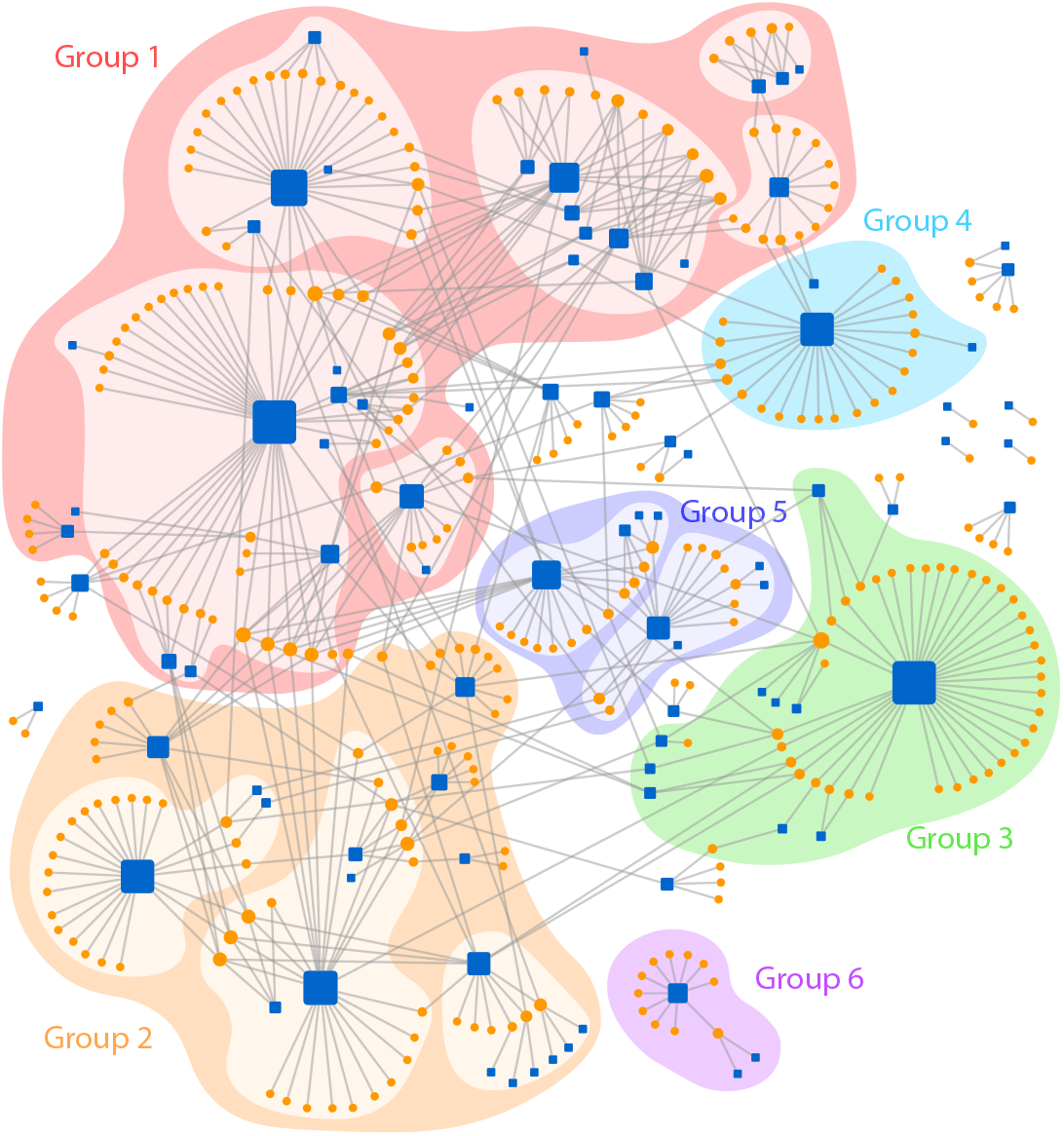
Territories in the odorant-receptor network. The six largest global groups, ranked by the number of receptors in the group, are shown in distinct colors. The sub-structures (local groups), if any, are also shown in lighter shades.

### Odorants in the same group tend to carry similar olfactory features

Our analysis so far only concerned the receptor codes for the test odorants; we did not use any other information about the odorants. On the other hand, because the dataset was constructed to include those odorants that were considered in previous psychology studies [13, 14], each test odorant has a known perceptual quality; i.e., how it smells to human.

It turns out that the two hub odorants in the largest receptor-code-based group (Group 1) represent the two sub-classes of plant-related odors: the characteristic “green” odor of fresh-cut grass (cis-3-hexen-1-ol) and the characteristic “floral” odor that is encountered in many natural essential oils (geranyl acetate). Moreover, there are many other odorants in Group 1 that belong to similar perceptual categories, namely green (phenyl acetaldehyde, sandalwood, terpineol); floral (linalool, lilial, lyral, anisaldehyde); and minty (methyl salicylate, spearmint, and the carvones). In Group 2, all three hub odorants are related to the body odor. Androstenone and androstadienone (the “human pheromone”) are contained in sweat or saliva; butyric acid with its characteristic sweaty smell is naturally formed in the human body by fermentation, and is also found in milk and butter. Among the smaller-degree odorants, heptanoic acid carries a rancid sweaty odor, and 2-methyl-1-propanethiol is associated with meaty or beer-like flavors. In Groups 3, 4 and 5, we find the ingredients of culinary spices. The two hub odorants in Groups 3 and 4 are the major constituents of clove oil (eugenol and eugenol acetate); the two hub odorants in Group 5 are common food-flavoring agents (ethyl vanillin and cinnamon). Group 3 includes more spice- or food-related odorants: nutmeg and banana have their well-known flavors, guaiacol is known to give the characteristic flavor to whiskey and roasted coffee, and 2-heptanone is related to the typical smell of gorgonzola cheese.

These observations lead to a hypothesis that, whereas our analysis is based only on the receptor codes and their redundancy, it results in a grouping of perceptually similar odorants together. In order to make the idea more quantitative, we labeled each odorant with a coarse-grained *olfactory feature* out of five categories (plantlike, animal-like, culinary, citrus, and others). More specifically, we first labeled the odorants using the eight perceptual odor categories identified from a previous work [21] (the primary odor category), then merged some of the categories that show similar co-occurrence patterns as described below (the secondary odor category); also see Fig. S7c and Table S1. Note that throughout this paper, we will use the term “olfactory features” specifically to refer to the perceptual odor qualities that are labeled by a discrete set of categories. Details of the labeling procedures are described in Methods. As a result, each odorant is assigned two independently determined labels: one for the receptor-code group (the “group”) and another for the perceptual odor category (the olfactory “feature”). Similarly as with the receptor-code groups, in this analysis we will consider the secondary odor categories which provide a more parsimonious representation of the perceptual odor space.

We sampled the joint distribution of receptor-code groups and olfactory features in the 89 test odorants, where each odorant is weighted by its degree so that its impact on the receptor space is taken into account (see Methods). We can say that an olfactory feature is *enriched* in a given group if the pair of two labels is observed more frequently in the joint distribution than expected from the marginal distributions of the individual labels. In particular, we quantified their joint enrichment in terms of the weighted log likelihood ratio (Fig. 6a), which sum up to give the mutual information between the two labels (see Methods). We find strong enrichments of plant-like odor in Group 1, animal-like odor in Groups 2 and 6, and culinary (food-related) odor in Groups 3, 4 and 5, supporting the hypothesis stated above. We also find that citrus is enriched in the union of all smaller groups, with the size-ranked group indices 7 and above. We performed the G-test and the chi-squared test to evaluate the statistical significance of the claim, and obtained *p* < 0.001 from both methods (*G* = 57.5 and χ^2^ = 62.2 for 24 degrees of freedom; see Methods for details).

**Figure 6:**
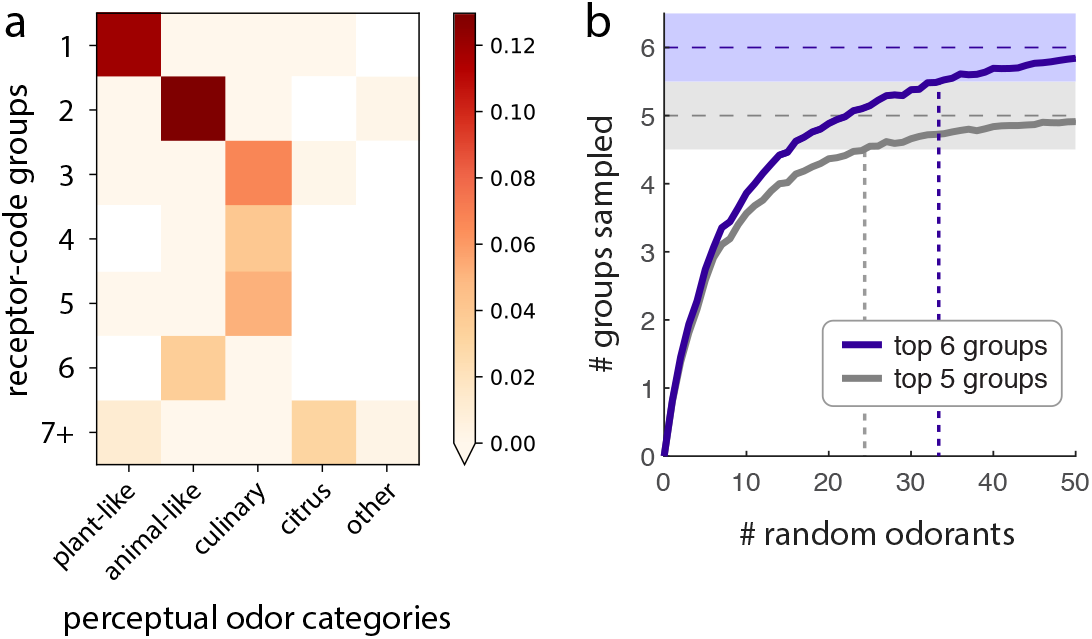
Link to the perceptual odor categories. **(a)** The joint enrichment between the receptor-code groups and the perceptual odor labels, measured in terms of the weighted log likelihood ratios. Also see Fig. S7. **(b)** The average number of major groups, out of six, that are covered by a random mixture of odorants as the number of odorants is varied (purple). The group coverage was closer to 6 than to 5 at *m* ≳ 33. Result is based on 500 independent sampling for each *m*, and the standard errors are smaller than the line width. If we instead consider up to five groups, the cutoff is *m* ≳ 25 (gray).

### Groups in the odor space explain the “olfactory white”

Lastly, we show that our results can be connected to the reported phenomenon of “olfactory white” [22], in which human subjects cannot distinguish odor mixtures if more than 30–40 odorants are randomly mixed while spanning the odor space. However, the idea of “spanning the odor space” remains at the level of a qualitative analogy to lower-dimensional sensory modalities such as color vision or tonal audition. In this study, we showed that the olfactory space can be understood in terms of a small number of (six) groups, both at the level of receptor codes and of perceptual features. Now we explore the idea of olfactory white more quantitatively, by asking when the odor space, partitioned in terms of the six major groups identified from the receptor codes, is fully covered by a mixture of odorants.

We sample a random set of *m* odorants from our test dataset, and count the number of major groups (out of six) with at least one odorant in the sampled set. Repeating the sampling while varying the mixture size *m*, we examine the average number of groups covered by the mixture (Fig. 6b). It turns out that a random sampling of *m* ~ 33 odorants is just enough to sample each of the 6 major groups at least once, in the sense that the average group coverage is > 5.5. On the other hand, the average receptor coverage (the number of receptors that recognize at least one odorant in the mixture) by a mixture of *m* ~ 34 random odorants is equivalent to the coverage by the 6 hub odorants (Fig. S3b). Therefore, we can say that a random mixture of 30–40 odorants is equivalent to a complete spanning of the odor space (up to the six major groups), in terms of the group coverage (number of bases spanned) as well as in terms of the receptor coverage. Our characterization of the human olfactory space, in terms of the receptor-code groups, offers a quantitative account for the number of random odorants needed to construct perceptually convergent odor mixtures (“olfactory whites”).

## DISCUSSION

We discovered a natural partitioning of the receptor code, by identifying groups of olfactory receptors (ORs) with correlated response patterns. The same procedure also identifies groups of odorants with similar receptor codes. Whereas the grouping was performed without any external label other than the receptor codes, we find that odorants in the same group tend to carry similar olfactory features, which indicates the existence of “labeled groups” in the receptor code space. Below we will elaborate this idea and discuss its implications from a broader perspective of human odor perception.

### Receptor groups as the bases of perceptual odor space

The presence of perceptually labeled receptor groups implies a lowdimensional nature of olfactory processing. Because the odor space is represented by the response pattern of *N* ≈ 400 functional ORs, there are in principle 2^400^ possible OR states, even in the binary regime. But when a group of receptors respond in a correlated way, not all 2^400^ states are equally likely. Therefore, a significant amount of dimensionality reduction is made in the receptor code space. This aligns with the previous ideas involving the effective sparsity of odor space that, despite the apparent high-dimensionality implicated by physiochemical properties of odorants, olfaction is working in a much lower dimension in effect [23–25]; or that the odor space is intrinsically clustered rather than uniform [21, 22].

The effective sparsity of the receptor code space, which may once again propagate to the perceptual odor space, has an analogy to vision. When one says that the natural visual scene is sparse [26], it means that the scene consists of a small number of features, such as lines or edges [27]. Here we take a viewpoint that the coarsegrained *olfactory features* (labeled by a small number of odor categories) are the “lines and edges” of the perceptual odor space, and that receptor groups represent the receptive fields for these features, with correlated response patterns. Whereas feature-extracting coding was thought to be the realm of higher-level neurons [28, 29], here we found a concrete evidence that it already starts at the level of receptors. It remains an open question to study how the encoded information is further organized through the downstream interaction between the OR neurons [30, 31].

### Hard-coded labels in the receptor space

Our findings of labeled groups support the idea that primary features of olfactory perception may already be “hard-coded” in the receptor code space [32, 33], enabling innate as well as learned behavioral responses. Indeed, if there exist some basic olfactory features relevant to the core functions of the organism, there is a clear advantage in keeping hard-coded labels for such features: fewer layers would enable a faster information processing [34] that does not require higher-level neural computations. It is also likely that such important features are the most strongly conserved over the course of evolution of the olfactory genome [35]. In fact, most examples of labeled-line receptors in other organisms are coarse-grained discriminators for certain groups of odorants [36, 37], or simply for good versus bad [33, 38], which is arguably the most important axis of odor perception for humans as well [24, 39, 40]. This extends the idea of *labeled lines*, in the sense that a group of receptors maintains a set of dedicated channels for a distinct perceptual feature.

However, we also note that the labeled-lines picture is not a perfect explanation of the data. The grouping of receptor codes is itself noisy (there are false positives and false negatives in the co-activation matrix of receptors), and the perceptual labeling of groups is noisy as well (e.g., there are odorants that belong to Group 2 yet carry plant-like odors). This implies that the receptor code also has signatures of *combinatorial coding*, a flexible point of view that assumes that the collective activities of receptors (rather than the activation of any specific receptor) encodes the odor information. For example, odorants that belong to the same receptor-code groups may carry different perceptual meanings (e.g., whereas all receptors that respond to the odorant 1-octanethiol also respond to cinnamaldehyde, 1-octanethiol smells like sulfur while cinnamaldehyde carries a pleasant, cinnamon-like scent). Our results therefore represent a hybrid between the two coding strategies. This adds to the growing evidence that olfactory coding in most organisms, including humans, are clearly more complicated than the dichotomy between labeled lines and combinatorial coding [41–43], although there are some cases in which one strategy seems to dominate [4, 38].

### More on the importance of hub odorants

At the core of our analysis are the hub odorants that elicit responses in the large population of receptors. The hub odorants create an important structure in the receptor space. According to our statistical analysis, the receptor codes for the largest hubs are almost non-redundant, and they “span” the receptor code space. If we take seriously the proposal that structures in the receptor code space represent the features in the perceptual odor space, we may further hypothesize that the hub odorants correspond to the important stimuli in the natural olfactory environment.

On the other hand, the six hub odorants in the dataset are described as the “typical” or “characteristic” odor of certain category, which are also commonly encountered in a human’s life. For example, cis-3-hexen-1-ol is recognized as the characteristic grassy smell; geranyl acetate is found in many natural essential oils and carries a floral aroma; butyl acetate and androstenone are found in the human and animal body; eugenol (clove), cinnamon, and ethyl vanillin (vanilla) each represents a characteristic and widely used ingredient for food flavoring.

Taking the observations together, we hypothesize that the receptor code is designed to assign a larger number of receptors to a more *salient* stimulus, whose presence is more strongly correlated with a particular olfactory feature. The existence of salient odorants in the natural odor space is in fact plausible from the biological viewpoint. For example, because the types of odorants produced by the flower is an outcome of evolutionary selection, most flowers have a few common odorants, with their physiochemical properties attracting the pollinating insects [44]. When more salient stimulus is encoded by the response of more ORs, it has the advantage that important olfactory signals (carried by the salient odorants) are detected reliably: i.e., responses are not lost under perturbations in the receptor space, such as the functional loss of certain ORs due to genetic mutation [45, 46]. If the hypothesis was correct, one would be able to predict the saliency of odorants in the natural odor space based on the number of receptors that respond. This opens up a series of problems related to the natural odor statistics. In particular, an interesting problem would be to explore whether the statistical salience has any relationship with the statistics of co-occurrence between odors [47].

### Limitations and open problems

Here we briefly discuss the limitations of our current approach, and the related problems for future research.

First of all, our analysis considers a binary approximation of the receptor code space, marking an odorant-receptor pair as “interacting” if there is a response at all. The ultimate understanding of the receptor codes should also incorporate the higher-order effects, such as the dependence on odor concentration [17, 48]; the effect of inverse agonists [49]; or the interaction between different odorants [50]. Only then will it be possible to fully address how the information in the receptor code is collectively processed, by the inter-glomerular interactions [30, 31]; with the temporal effects [51] involving sniffs [52]; or under different abundances of the receptors [53]. As for the concentration dependence, we show that the current framework can be extended by constructing a family of binary interaction networks at varying odor concentrations (Fig. S2). The effect of odorant concentration on receptor code is in principle straightforward to analyze under our odorant-receptor based framework even for a mixture of odorants at heterogeneous concentrations; however, meaningful investigation of such effect on human olfactory sensing would require a more systematic normalization with respect to the human detection thresholds that are different for the individual odorants [40, 54]. Our work lays down a general framework on which more specific questions like this can be addressed.

Second, in light of the three levels of olfactory processing that we laid out at the beginning (Fig. 1), it is natural to expect that the receptor code space also reflects the chemical properties of the odorants that comprise the molecular space. Although we did not extend our analysis to the molecular space in this work, previous works suggest a correlation between the molecular similarity between pairs of odorants and the similarity of their olfactory responses at the level of OR neurons [12, 17, 48, 55]. Now that we have a low-dimensional characterization of the receptor code space in terms of the groups, an interesting problem would be to investigate the correlation between the receptor-code groups and the chemical properties of the odorants.

Third, whereas our analysis achieved an unprecedented link between the receptor code space and the perceptual odor space, further progress can be made a better characterization of the perceptual space. In particular, we relied on a specific mapping of the test descriptors to the set of reference words that describe the eight perceptual odor categories [21]. This process can be replaced by a more systematic quantification of similarity between the descriptor words, which would require the understanding of the underlying semantic space for olfactory descriptors.

### Concluding remarks

A set of concrete messages can be drawn from our study: (i) The existence of odorant groups, or the correlated patterns between the receptor codes, greatly reduces the dimensionality of the receptor code space and allows us to identify coarse-grained olfactory features in the perceptual odor space. (ii) The odorant and receptor groups revealed from our study clarify the balance between the labeled lines and combinatorial coding strategies. The labeled receptor groups on top of non-uniformly distributed receptor codes imply a hybrid strategy, which can be used to leverage both the fidelity of the labeled-line design and discriminative capacity of the combinatorial coding. (iii) Each large group of odorants is described by the coarse-grained olfactory feature (e.g. plant-related, animal-related odor).

In summary, by analyzing the dataset of OR responses, we identified a modular structure inherent in the receptor code space of human olfaction. Insights from these findings could be extended to the broader study of human odor perception and help understand exceptionally diverse data on olfaction in a more systematic manner.

## METHODS

### Modeling

#### Network representation of the receptor code space

We analyze the dose-response curves of 535 interacting odorant-receptor pairs, involving 89 odorants and 303 olfactory receptors (ORs), from the dataset of [14]. The interacting odorant-OR pairs were screened from a library of 511 human OR genes [13, 14]; the final number of 303 ORs include only those that respond to at least one odorant in the dataset. The panel of odorants was selected to span the 20-dimensional physiochemical space that explains the variance in mammalian OR responses [12]. The receptors in the dataset, which cover a near-full (~ 3/4) space of known human ORs, can be viewed as the receptor-space representation of the odorant. On the other hand, the set of cognate odorants found for each receptor, which is only a small sampling of the olfactory environment (humans can detect at least ≳ 10^4^ odorants), is not guaranteed to be complete. We note that the dataset also includes several natural odors that are not monomolecular odorants (for example “banana”), which do not exactly fit in our formulation of pairwise odorant-OR interactions. Nevertheless, these are only a minor fraction of the dataset, and we included all reported odors in our analysis.

A graph can be formally written as *G* = (*V, E*), where *V* is the set of all nodes (“vertices”) in the graph, and *E* is the set of all edges. In the binary regime, each edge in *E* is an unordered pair of nodes in *V*; this gives a binary graph, which can also be represented in terms of an *adjacency matrix A*, whose (*i, j*)-th element *A_ij_* represents whether there is an edge between the two nodes *i* and *j* in the graph. Specifically, the odorant-receptor network is a *bipartite graph*, where each node belongs to either of two node types (either odorants or receptors) and every edge connects one odorant node to one receptor node. If *V_O_* and *V_R_* are the sets of all odorants nodes and the set of all receptors nodes, respectively, each edge in *E* connects exactly one node in *V_O_* and the other node in *V_R_*. This bipartite structure is cast into the *M* × *N* sub-matrix of the full adjacency matrix, *A*_λ*i*_, where *N* = |*V_R_*| is the number of receptor nodes in the network, and *M* = |*V_O_*| the number of odorant nodes. Each element of the matrix is defined as *A*_λ*i*_ = 1 if and only if there is an edge between odorant *λ* and receptor *i*. We call *A*_λ*i*_ the *interaction matrix*.

#### Binary vector representation of a receptor code

The receptor code space representation of an odorant, written in a binary vector **y**, is equivalent to the corresponding row of the interaction matrix. That is, [**y**_λ_]_*i*_ = *A*_λ*i*_, where [*y*]_*i*_ is the *i*-th element of vector **y**.

The union of two binary vectors is defined element-wise as [**y**_1_ ∪ **y**_2_]_*i*_ = [**y**_1_]_*i*_ ∨ [**y**_2_]_*i*_, where ∨ is the logical “or” operation between two binary variables. Similarly, the intersection is defined as [**y**_1_ ∩ **y**_2_]_*i*_ = [**y**_1_]_*i*_ ∧ [**y**_2_]_*i*_, where Λ is the logical “and” operation. We also define the difference of two binary vectors as [**y**_1_ \ **y**_2_]_*i*_ = [**y**_1_]_*i*_ ∧ ¬[**y**_2_]_*i*_, where ¬ is the logical “not” operation. For the union and the intersection, the definitions easily extend to more than two binary vectors.

#### Degree of an odorant

The *degree* of a node is the number of edges attached to it, or (when there is no loop) the number of other nodes that are connected to it. It is useful to define the *size* of a binary vector as the number of its non-zero elements: 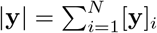.

The degree of an odorant node is the number of receptor nodes connected to it:

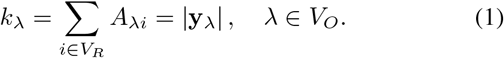

In the current study, we primarily consider the odorant degree, or how broadly (or narrowly) an odorant is “targeting” the receptors. This breadth of interaction for the odorant is well-defined, because there is a finite number of olfactory receptors. The dataset in [14] covers a majority of the known (functional) human olfactory receptor repertoire: 303 out of ~ 400 ORs. Our approach has advantage over other studies that considered whether a given receptor is “narrowly tuned” or “broadly tuned” across different odors, which is hard to quantify without a good metric of the odor space.

The degree distribution of odorants, *P_O_*(*k*), is measured by counting the number of odorants at each degree *k*. *P_O_*(*k*) appears approximately straight in the log-log scale, and is fitted to the curve, *P*(*k*) ~ *k*^−*α*^. However, the fit does not necessarily indicate an underlying distribution that is power-law in a strict sense; we are simply using the power-law dependence to describe the heavy-tailed shape of the degree distribution.

#### Co-activation graph

The *co-activation graph* is a projection of the original bipartite graph to one of the two node types. For example, the co-activation graph of receptors, *G_R_* = (*V_R_, E_R;O_*), inherits all receptor nodes *V_R_* from the original bipartite graph. Two receptor nodes in the co-activation graph *G_R_* are connected by an edge if there is a common odorant node in the original graph *G* that is connected simultaneously to both receptors (see Fig. S4a,f). To define the edges, *E_R;O_*, it is sufficient to define the corresponding adjacency matrix *A_R_*, where (*A_R_*)_*ij*_ = 1 indicates an edge between two receptors *i, j* in the co-activation graph. We call *A_R_* the *co-activation matrix* of receptors. Each element of the co-activation matrix is given as (*A_R_*)_*ij*_ = ∨_λ∈*V_O_*_ (*A*_λ*i*_ Λ *A*_λ*j*_), which indicates whether there is at least one odorant λ for which *A*_λ*i*_ and *A*_λ*j*_ are both 1, where *A*_λ*i*_ is the interaction matrix for the original bipartite graph.

### Receptor code redundancy

We define the receptor code redundancy as χ = (*n*_full_ − *n*_eff_)/*n*_full_, where *n*_full_ is the full impact of a signal in the absence of redundancy, and *n*_eff_ is the net effect of the signal under redundancy.

#### Receptor code redundancy of a target with respect to the background

For a discrimination task of a target odorant λ in the presence of a background odorant *μ*, the effect of redundancy reduces the net effect of adding λ from *n*_full_ = |**y**_λ_| to *n*_eff_ = |**y**_λ_ \ **y**_*μ*_|. In this case, the receptor code redundancy is χ_λ_|_*μ*_ = (|**y**_λ_| − |**y**_λ_ \ **y**_*μ*_|)/ |**y**_λ_| = |**y**_λ_ ∩ **y**_*μ*_| / |**y**_λ_|.

#### Receptor coverage by a mixture of odorants

We can also consider an odor signal that consists of a mixture of odorants. We define its *receptor coverage* as the number of receptors that respond to at least one odorant in the mixture, *n*_union_ = |**y**_1_ ∪ ⋯ ∪ **y**_*m*_|. We can also count the sum of individual odorant degrees, *n*_sum_ = |**y**_1_| + ⋯ + |**y**_*m*_|. Note that the net receptor coverage *n*_union_ corresponds to *n*_eff_, and the sum of individual coverages *n*_sum_ to *n*_full_, in the above construction (also see Fig. S3a).

#### Receptor code redundancy in a mixture of odorants

The two numbers, *n*_union_ and *n*_sum_, are different when there are overlaps between the receptor codes for the odorants in the mixture. Therefore, we characterize the net amount of receptor code overlap as *n*_sum_ − *n*_union_, and define the receptor code redundancy in a mixture as χ_mix_ = (*n*_sum_ *n*_union_)/*n*_sum_.

#### Odorant mixture test

We selected mixtures of *m* odorants from the interaction network, either randomly (random mixture), or selecting the highest-degree odorants first (mixture of hubs). For each mixture, we counted the receptor coverage *n*_union_ and the sum of individual coverages *n*_sum_. For the random mixtures, we drew 500 independent samplings at each *m*; because we tried 90 different values of 0 ≤ *m* ≤ 89, we have 45,000 random mixture samples.

The receptor coverage by the six hub odorants is *n*_union_ = 155, about half of the receptor space (Fig. S3b). However, its receptor code redundancy is only χ_mix_ = 0.08. This is much smaller than what is expected from a random-odorant mixture at the same receptor coverage (χ_mix_ ≈ 0.23 using the average counts); none of 105 random mixtures at the fixed receptor coverage *n*_union_ = 155 had a smaller redundancy (*p* < 0.01). Considering all mixtures with *n*_union_ ≤ 155, we got *p* < 0.02 (317 out of 17340). If we consider all random mixtures and count the samples that are below the red curve in Fig. S3c, we get *p* = 0.05 (2420 out of 45,000), where we note that most of the extreme samples come from the top end of the curves, where the newly added odorants are no longer “hubs”. Besides, the average redundancy curves show that the level of redundancy with the six hubs (χ_mix_ ≈ 0.1) is reached already at a receptor coverage of *n*_union_ ≈ 70 in an average random mixture, which corresponds to *m* ≈ 13 odorants (Fig. S3c).

We also note that the redundancy curve for the mixture of hubs (Fig. S3c, red) deviates most greatly from the “chance curve” obtained from the random mixtures (gray) when the six hub odorants were selected. This adds support to our focus on the six hub odorants.

### Grouping in the receptor code space

We will provide details of the grouping process in three steps: node ranking, primary grouping and secondary grouping. To start, we rank the odorants and receptors in the dataset.

#### Ranking odorants and receptors by the degrees

Odorants are ranked by their degree of interaction, *k* (Eq. 1). When there are multiple odorants with the same degree, they are ranked according to the order they are indexed in the original dataset [14]. We call this ordering *h*_0_, such that each odorant λ (where λ may be an arbitrary index) is assigned a unique rank *h*_0_(λ) ∈ {1, ⋯, *M*}.

Receptors are ranked by their degree in the co-activation graph. The degree in the co-activation graph represents the number of other receptors that share at least one odorant partner with the given receptor. When there are multiple receptors with the same degree of co-activation, they are ranked according to the order they are indexed in the original dataset [14]. Similarly as for the odorants, we call this ordering *g*_0_, such that each receptor *i* is assigned a unique rank *g*_0_(*i*) ∈ {1, ⋯, *N*}.

#### Primary grouping

We first partition the receptor space into a set of non-overlapping groups, and subsequently use these to classify the odorants based on the receptor code overlap.

In the primary receptor grouping (*g*_1_), receptors are grouped such that each receptor “signs up” to the largest receptor clique it belongs to. Specifically, we assign to each receptor the rank *h*_0_ of the highest-degree odorant it interacts with; in case of ties, we choose odorants of higher ranks (smaller *h*_0_). We re-rank the receptor group index while preserving the order of the associated rank *h*_0_, by assigning the primary group index *I* = 1 to the largest receptor group, and the index *I* = 2 to the next largest group, and so on (*I* =1, 2, ⋯ *I*_max_). This results in a grouping *g*_1_: *i* ↦ *I* that maps each receptor *i* to a group index *g*_1_(*i*) = *I*.

In the primary odorant grouping (*h*_1_), each odorant is assigned to the above-determined receptor group (*g*_1_) to which its receptor code overlaps the most. The primary odorant grouping *h*_1_: λ ↦ *I* (or *h*_1_(λ) = *I*) is defined by choosing the receptor group of index *I*, with which the odorant’s receptor code (**y**_λ_) displays the maximum overlap, namely *I* = argmax_J_ χ_λ|J_. Here, the receptor code overlap of an odorant λ to a receptor group index *I* is quantified as χ_λ|*I*_ = |**y**_λ_ ∩ **y**_*I*_| / |**y**_λ_|, where the binary vector **y**_*I*_ represents the receptor code for the group *I* by having each element (**y**_*I*_)_*i*_ = 1 if *g*_1_(*i*) = *I*, and (**y**_*I*_)_*i*_ = 0 otherwise. Note that the resulting odorant group indices span through *I* =1,2, ⋯, *I*_max_, shared with the receptor group indices.

#### Secondary grouping

We merge the primary groups based on the amount of co-activation in the receptor code space.

In the secondary receptor grouping (*g*_2_), pairs of primary receptor groups are merged if there is a strong co-activation. For two primary receptor groups *I*, *J*, we calculate the amount of coactivation between the two groups, simply in terms of the density of off-diagonal-block contribution in the co-activation matrix *A_R_*:

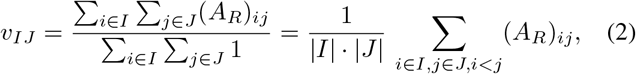

where *i* ∈ *I* is a shorthand to indicate that the sum runs over all receptors *i* such that *g*_1_(*i*) = *I*. We merge each pair of primary groups *I* and *J* whenever their co-activation is stronger than a threshold, i.e., if 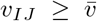. Here we used an optimal threshold value of 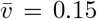, where the grouping was most informative (Fig. S5; also see Statistical validation). Note that this is a greedy and transitive grouping. For example, if there are three primary groups *I*_1_, *I*_2_, *I*_3_ such that 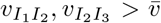 but 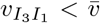, then all three are merged together in the secondary grouping; the two groups *I*_1_, *I*_2_ are first combined, then *I*_3_ is also pulled into the group that *I*_2_ belongs to. After performing all pairwise merges, we once again re-rank the resulting group indices while preserving the order. This defines a grouping *G*_12_: *I* ↦ *I*′ that maps each primary group *I* to a (re-numbered) secondary group *I*′. Each receptor *i* is then assigned a secondary group index *g*_2_(*i*) = *G*_12_(*g*_1_(*i*)).

In the secondary odorant grouping (*h*_2_), pairs of primary odorant groups are merged if there is a strong co-activation. Because the primary grouping was a simultaneous partitioning of odorants and receptors, the above merging operation of receptor groups from *g*_1_(*i*) to *g*_2_(*i*) automatically propagates to the odorant groups. Specifically, each odorant λ is assigned a secondary group index *h*_2_(λ) = *G*_12_(*h*_1_(λ)).

### Labeling with the perceptual odor categories

#### Descriptor space

The perceptual odor quality of each odorant is characterized by a list of odor descriptors (Table S2). We collected the odor descriptors for the 89 test odorants in the dataset [14], from multiple sources including public databases, academic works and industrial reports (see Table S5 for data sources). In particular, we obtained a majority of odor descriptors from the Good Scents Company Information System [56], using custom scripts for web-scraping.

Let *W* be the set of all descriptors that appear at least once in our collection. We have *N_W_* = |*W*| = 194 unique descriptor words for the *M* = 89 test odorants in the dataset [14]. We can think of an *M* × *N_W_* binary matrix, 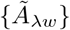, such that 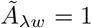 if odorant λ is described by the word *w*, and 0 otherwise.

#### Perceptual odor categories

We adopt the 8 odor categories identified in a previous work [21]. Each category *s* ∈ {1,2, ⋯, 8} is defined by a list of reference descriptors, 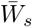, provided in Figure 7 in the original paper (also shown in Table S3). For convenience, we also nickname each category with a single representative descriptor: (1) citrus, (2) chemical, (3) minty, (4) floral, (5) fruity, (6) green, (7) spicy, and (8) animal. The representative nickname for each category was chosen according to our own semantic senses; its only role is to provide an intuitive name by which we can easily refer to the category, and it does not play any further role in the analysis.

#### Labeling odor descriptors

We first label each descriptor *w* ∈ *W* with a unique perceptual category *s* ∈ {1, 2, ⋯, 8, 9}, where the additional 9th category stands for “unsorted”. This defines a map 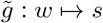. Here we only consider a deterministic categorization of descriptors, such that 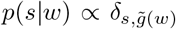. The mapping is straightforward when the descriptor is directly contained in the reference vocabulary; because 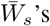 are non-overlapping by construction, we can find a unique category *s* for the descriptor in this case. However, some descriptors do not exactly match with any of the reference words, in which case the mapping requires a sense of similarity between the words in the underlying semantic space.

Here we attempt to make a projection of each descriptor onto the space of reference words. Specifically, for each test descriptor *w*, we choose a proxy 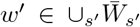 from the reference vocabulary that can approximate the descriptor *w*, and identify the category s for which 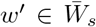. The extra category “unsorted” is assigned to abstract or ambiguous descriptors [57]. Null descriptors like “odorless” or “weak” are also labeled as unsorted. The proxies were determined manually by the authors using the human sense of the semantic space, and the entire mapping is made available as a supplementary table (Table S4) so that the reader can reproduce and/or judge the procedure.

#### Labeling odorants

Next, we label each odorant λ with a perceptual category *s*. To summarize how each odorant is associated with a set of descriptors, we write a conditional probability

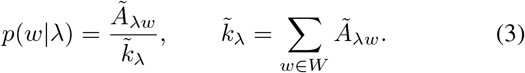

This allows us to write *p*(*s*|λ) = Σ_*w∈W*_ *p*(*s*′|*w*)*p*(*w*|λ). For simplicity, we consider a deterministic map that involves a “vote” by all descriptors of each odorant:

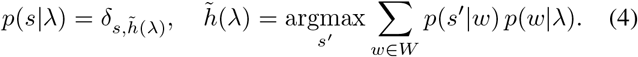

In case of ties, we put more weight to the descriptor that was marked as the representative “odor type” in the GoodScents database [56]. Now we have a map 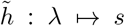 that assigns a perceptual odor category *s* to each odorant λ.

#### Joint distribution of receptor-code groups and perceptual odor categories

Treating the receptor-code group index *s_R_* and the perceptual odor category index *s_W_* as random variables, we can formally write down their joint probability distribution as:

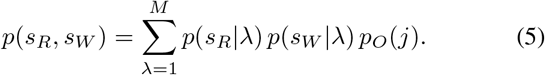

This is to say that the two group/category indices are *sampled* by the test odorants, where the two grouping maps *p*(*s_R_*|λ) and *p*(*s_W_*|λ) are conditionally independent for a given odorant λ.

Because we already defined the two deterministic maps, we can write down the two conditional distributions as *p*(*s_R_*|λ) = *δ*_*s*_*R*_,*h*(λ)_ and 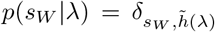. The third term, *p_O_*(λ), represents the weight to each odorant. If we were to count all odorants equally, we would have used a uniform *p_O_*(λ) = 1/*M*. However, we would like a sampling that is aware of the odorant’s “weight’ in the receptor space, such that an odorant that affects more receptors (a higher-degree odorant) counts more. To this end, we write

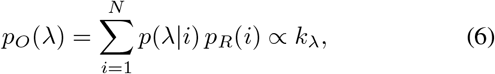

where 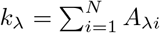 is the degree of the odorant λ in the interaction network. Note that this approach assumes a uniform *p_R_*(*i*) ∝ 1 for the receptors, rather than the odorants. In this case, *p_O_*(λ) can be interpreted as the probability that a randomly chosen receptor recognizes the odorant λ. Summarizing, the joint distribution is sampled as

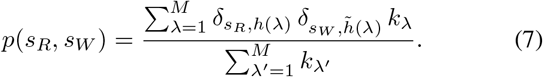

We can also consider the two marginal distributions, *p*(*s_R_*) = Σ_*s_W_*_ *p*(*s_R_, s_W_*) and *p*(*s_W_*) = Σ*_s_R__ p*(*s_R_, s_W_*), and consequently the two conditional distributions, *p*(*s_R_|s_W_*) ≡ *p*(*s_R_, s_W_*)/*p*(*s_W_*) and *p*(*s_W_|s_R_*) ≡ *p*(*s_R_, s_W_*)/*p*(*s_R_*).

### Statistical validation

#### Optimal threshold for secondary receptor grouping

In the secondary receptor grouping, our goal was to obtain the best partitioning of the receptors so that the resulting structure captures the co-activation pattern. Because we start from the primary receptor groups as the “units”, and work by merging them based on the overlaps, the grouping depends on the threshold 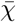. In the limit where the threshold is too small, most units are merged together and a giant cluster is formed (Fig. S5a). On the other hand, when the threshold is too large, most units stay un-merged, failing to capture the global structure (Fig. S5c). The optimal threshold is where the grouping is most *informative* (see below), in the sense that more strongly co-activating primary groups are merged together in the same secondary group, and non-co-activating groups stay apart (Fig. S5b).

Here we consider our grouping as a binary classifier, where each receptor pair has one of the two real states (either co-activated by an odorant or not), and one of the two predicted states (either grouped together or not). We count the number of receptor pairs that belong to each of the four outcomes: co-activated and grouped (true positive; TP), co-activated and ungrouped (false negative; FN), non-co-activated and grouped (false positive; FP), and non-co-activated and ungrouped (true negative; TN). We also define the real positive (RP = TP + FN) and the real negative (RN = TN + FP) in the data, and similarly the predictive positive (PP = TP + FP) and the predicted negative (PN = TN + FN) according to the grouping. There are multiple ways to quantify the informativeness of the classifier, but the basic idea is to understand the tradeoff between the type I (FP) and type II (FN) errors. Here we take two common approaches, considering the sensitivity-specificity and the precision-recall tradeoff respectively. The *sensitivity* is defined as the true positive rate TPR = TP/RP, which is also called the *recall*. The *specificity* is defined as the true negative rate TNR = TP/RN. It is also common to write the specificity in terms of the *fall-out*, or the false positive rate FPR = TP/RN with a simple relationship FPR =1 - TNR. The *precision* is defined as the positive predicted value PPV = TP/PP. Also see Fig. S5d.

To find the optimal grouping in terms of the sensitivity and the specificity, we minimize the sum of squared error, SSE = FNR^2^ + FPR^2^ = ((1 - sensitivity)^2^ + (1 − specificity)^2^). This is also graphically equivalent to finding the point that is closest to the upper left corner of the receiver operating characteristic (ROC) plane, which plots the sensitivity versus the 1-specificity at varying threshold 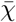 (Fig. S5e-f).

Alternatively, we can use the *F_β_* score, another popular quantity defined as the weighted harmonic average of precision and recall [58]:

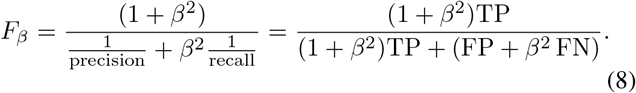

A larger *β* puts more weight to the recall (pays more attention to the false negatives), whereas a smaller *β* puts more weight to precision (more attention to the false positives). Because the actual dataset is strongly imbalanced such that RN > RP, we choose a value of *β* that penalizes false negatives more heavily [59]. Specifically, we use 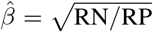, so that the weighted combination becomes 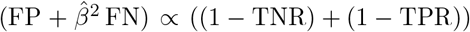. We can also plot the precision-recall curve as the threshold is varied (Fig. S5h-g). In our case, the two optimization conditions (minimum SSE and maximum *F_β_*) are satisfied at the same threshold 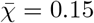, which defines the best secondary receptor grouping (Fig. S5f,h).

#### Shuffling test for receptor-code groups

To test the statistical significance of the observed receptor groups, we compare the result to three null models constructed by shuffling the odorant-receptor interactions. We consider three null models, with shuffling rules that preserve different marginal statistics.

First, we make a full shuffling of the activation matrix, *A*_λ*i*_, such that the number of non-zero elements (the number of interactions in the network) is preserved (Fig. S6a). This is also equivalent to fixing the average degree of the odorants.

Second, we construct a more constrained null model where the degree distribution is preserved. Specifically, we shuffle each odorant-row of the activation matrix *A*_λ*i*_ independently, such that each odorant re-connects to the receptors randomly, while the degree is kept fixed (Fig. S6b).

Third, to test the effect of the secondary grouping separately from the primary grouping, we consider a null model that preserves the primary grouping structure (Fig. S6c). For each odorant λ in a primary group *g* = *h*_1_(λ), We shuffle the the set of receptors that belong to the same primary group, {*i* ∈ {1, ⋯, *N*} | *g*_1_(*i*) = *g*}, and separately shuffle the rest of receptors that belong to all other primary groups, {*i* | *g*_1_ (*i*) ≠ *g*} (also see Fig. S6d). But as a result of this shuffling, the goodness of the original primary grouping becomes exaggerated, because shuffling destroys all residual correlation that was not captured by the primary groups. So we add extra randomness to restore the precision of primary grouping as it appears in the original data. We do this by making an appropriate number of *degree-preserving swaps*, as suggested by [60]. The degree-preserving swaps are made in the following way. We randomly choose 2 columns and 2 rows from the activation matrix, which defines a 2 × 2 binary sub-matrix. If the sub-matrix is either diagonal or anti-diagonal, we replace the sub-matrix with the other form, and count this as one swap. It is clear that such a swap preserves both the column-sum and the row-sum of the matrix. If the chosen sub-matrix has neither of the two forms, no action is made, and we proceed to sample a new sub-matrix. The two goodness measures are simultaneously restored to the levels that correspond to the original primary grouping, with 40 extra swaps (Fig. S6d).

Under each null model, we draw multiple shuffles to sample the distributions of the test statistics, in this case the two goodness measures SSE and *F_β_*. For each shuffle of the interaction data, we constructed the corresponding receptor co-activation matrix, and applied the grouping algorithm at varying overlap threshold 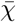. We chose the best grouping results as the test statistics for the sample, either at minimum SSE (Fig. S6e-f) or at maximum *F_β_* (Fig. S6g-h). The two measures were optimized separately, and may correspond to different optimal thresholds for the same shuffled sample. This procedure allows us to estimate the (one-tailed) p-value, which indicates the probability to have a grouping result from a shuffled dataset that is as good as or better than what we observe from real data. Because there was only up to 1 out of 500 shuffled samples that clusters better than the original dataset, we can say that *p* ≤ 0.002 (Fig. S6i-j).

#### Joint enrichment between receptor-code groups and perceptual odor categories

To quantify the correlation between the receptor-code groups *s_R_* and the perceptual odor categories *s_W_*, we consider the likelihood ratio *p*(*s_R_, s_W_*)/*p*(*s_R_*) *p*(*s_W_*). If this ratio is larger than 1 for a given (*s_R_, s_W_*) pair, it indicates an over-representation (enrichment) of the pair in the dataset, compared to the expectation from the null hypothesis that the two labels are completely uncorrelated (Fig. S7a). Note that the likelihood ratio can also be interpreted as the enrichment of one label given another, if re-written as *p*(*s_R_*|*s_W_*)/*p*(*s_R_*) or *p*(*s_W_|s_R_*)/*p*(*s_W_*).

The weighted sum of the log likelihood ratios is the mutual information:

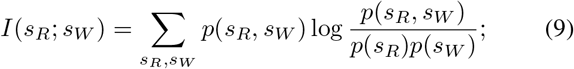

it is also equivalent to the Kullback-Leibler (KL) divergence, *D_KL_*[*p*(*s_R_, s_W_*)|||*p*(*s_R_*)*p*(*s_W_*)]. This sum quantifies the amount of information that is lost by incorrectly describing the observed joint distribution *p*(*s_R_, s_W_*) with the null hypothesis *p*(*s_R_*)*p*(*s_W_*).

It is also useful to look at each pair’s contribution to the mutual information, which has the form of a weighted log likelihood ratio *ρ*(*s_R_, s_W_*) = *p*(*s_R_, s_W_*)log[*p*(*s_R_, s_W_*)/*p*(*s_R_*)*p*(*s_W_*)]. For the purpose of understanding which pairs are more significantly enriched, we find it more informative to consider the weighted log likelihood ratio (Fig. S7b) than the log likelihood ratio alone (Fig. S7a). The weighting properly takes into account that the log likelihood adds up with independent observations. We will call *ρ*(*s_R_, s_W_*) the *joint enrichment* between the receptor-code groups *s_R_* and the perceptual odor categories *s_W_*.

#### Merging the categories for dimensionality reduction

Because the amount of information that can be drawn from data is fixed, it is important that we keep only as many groups/categories as relevant to our conclusion. In the receptor code space, we considered the six largest groups and treated the rest as a single extra group (6 degrees of freedom). In the perceptual odor space, we looked at which of the 8 odor categories have similar enrichment patterns (Fig. S7b), and also used the common-sense semantic knowledge, to merge the plant-like (green, floral, minty, chemical) and the culinary (spicy, fruity) labels together (Fig. S7c). The grouping of “chemical” with the plant-like category is based on the correlations in the enrichment pattern, although we should note that there is only one odorant in our dataset (quinoline) that is labeled as “chemical” according to our procedure. This results in a set of reduced perceptual odor categories with 4 degrees of freedom, labeled as plant-like, animal-like, culinary, citrus and others, which gives a more parsimonious classification of the perceptual odor space while preserving the enrichment pattern. For convenience, we distinguish the two set of labels with the names “Category 1” (the primary categories adopted from [21]) and “Category 2’ (the secondary categories after dimensionality reduction). The secondary category for the perceptual quality of an odorant is determined by its primary category (Table S1).

#### Evaluating the significance of group labeling

We used the G-test to evaluate the statistical significance of our finding that the receptor-code groups can be labeled with perceptual odor categories. The G-test statistic is defined as 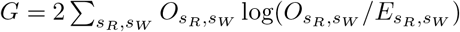, where *O_s_R_,s_W__* and *E_s_R_,s_W__* are the observed and the expected number of odorants with the pair of labels (*s_R_, s_W_*). By construction, we have *O_s_R_,s_W__* = *M_p_*(*s_R_, s_W_*) and *E_s_R_,s_W__* = *M_p_*(*s_R_*)*p*(*s_W_*), where *M* is the number of test odorants (observations). This gives *G* = 2*M* · *D_KL_*. We can calculate the p-value for this joint classification, by assuming that *G* approximately follows the chi-squared distribution with the appropriate degrees of freedom. In this case, the number of degrees of freedom for the chi-squared distribution is d.o.f. = (|*S_R_*| − 1)(|*S_W_*| − 1), where |*S_R_*| is the number of distinct receptor-code groups, and |*S_W_*| is the number of distinct odor categories. Using the primary odor categories, we get *G* = 68.9, which translates to *p* = 0.03 with 48 degrees of freedom (Fig. S7b). Using the secondary odor categories, we get *G* = 57.5 and *p* < 0.001 with 24 degrees of freedom (Fig. S7d).

To further validate our use of the G-test, we also compared the result with the chi-squared test, for which the test statistics is definedas 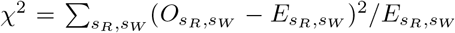. (Note that this χ^2^ should not be confused with the overlap measure χ or the overlap threshold 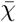 in our analysis of the receptor code space.) We get χ^2^ = 71.4 and *p* = 0.02 using the primary odor categories (Fig. S7e), and χ^2^ = 62.2 and *p* < 10^−4^ using the secondary categoreis (Fig. S7f). Both tests give small and consistent p-values, especially with the secondary odor categories with reduced degrees of freedom. This confirms the statistical significance of our findings, in which each major group in the receptor code space is associated with a characteristic label in the perceptual odor space.

## Data availability

The list of test odorants are provided in two sets of supplementary tables, with receptor-code groups and perceptual odor categories (Table S1) and with raw odor descriptors (Table S2) respectively. The list of reference words for each perceptual category [21] is provided as Table S3. Our labeling of each perceptual odor descriptor is shown in Table S4. Data sources are listed in Table S5.

Formatted data files to reproduce the network representation of odorant-receptor interaction with full attributes for each node and edge (as .cyjs and .cys files, to open in Cytoscape) are freely available at https://github.com/jihyunbak/ORnetwork.

## Code availability

The Matlab code for the grouping analysis, along with associated documentation, is available under the MIT license at https://github.com/jihyunbak/ORnetwork.

## ACKNOWLEDGMENTS

We thank the KIAS Center for Advanced Computation for providing computing resources. This study was partly supported by the grants from the National Research Foundation of Korea (NRF-2018R1A2B3001690) (C.H.), and the National Science Foundation (CHE-1362926) (S.J.J).

## AUTHOR CONTRIBUTIONS

J.H.B., S.J.J. and C.H. designed the projects. J.H.B. and C.H. performed research and analyzed the data. J.H.B., S.J.J. and C.H. wrote the paper.

## SUPPLEMENTARY MATERIALS

Supplementary Materials include seven figures and five tables, with accompanying text. All items are available at the end of this manuscript in the following order.

**Figure S1.** Interaction network with interaction properties.

**Figure S2.** Interaction network at fixed odorant concentrations.

**Figure S3.** Odorant mixture test.

**Figure S4.** Grouping procedure.

**Figure S5.** Finding the optimal threshold for partitioning.

**Figure S6.** Shuffling test for receptor code grouping.

**Figure S7.** Joint statistics between the receptor-code groups and the perceptual odor categories.

**Table S1.** List of odorants with receptor-code group and perceptual odor category labels.

**Table S2.** List of odorants with perceptual odor descriptors.

**Table S3.** Reference descriptors for the perceptual odor categories.

**Table S4.** Labeling odor descriptors.

**Table S5.** List of data sources.

## SUPPLEMENTARY TEXT

### Interaction network with full edge attributes

We start from previous characterization of the pairwise interactions in the dataset, based on a minimal kinetic model [61]. For each interacting pair of an odorant and a receptor, the dose-response curve *r*(*c|θ*), or the activity of the receptor in response to an odorant concentration c, is characterized with four parameters *θ* = {*B, E, K*_1/2_, *H*} as 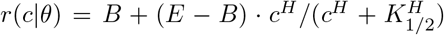. Here *B* is the basal activity of the receptor; *E* is the efficacy, or the maximum response from the interaction; *K*_1/2_ is the EC_50_, the effective concentration of the odorant required to elicit half-maximum response; and *H* is the Hill coefficient. The net amplitude (or the strength) of the response is measured by the difference *A* ≡ *E* − *B*. The interaction is either activating (*E > B*), or deactivating (*E < B*). In the previous study [16], we determined the four parameters {*B, E, K*_1/2_, *H*} for each of the 535 interacting odorant-receptor pairs from the dataset of [14]. The interaction parameters are displayed as the edge attributes in Fig. S1. A network file with full information on all nodes and edges, ready to import in an interactive platform (Cytoscape [62]), is also available online (https://github.com/jihyunbak/ORnetwork).

### Interaction network at fixed odorant concentrations

Here we show how the analysis used in our work can be extended beyond the binary regime by taking into account the effect of odorant concentration.

We construct a binary interaction network at a fixed odorant concentration, *c*, assumed to be uniform for all odorants considered. In this case, an edge between each odorant-receptor pair indicates that the receptor is *activated* when the odorant is present at this concentration *c*; this selects all activations (positive responses) with *c* > *K*_1/2_, and excludes the de-activations (negative responses). This binarization scheme is consistent with the prediction that the olfactory response is effectively binary at the level of receptor neurons [16]; it is also related to the ON/OFF model considered in [63]. Varying the odorant concentration *c* uniformly for all odorants, we obtain a family of binary (unweighted and undirected) interaction networks (see Fig. S2a). As expected, the average degree of an odorant decreases as c decreases (Fig. S2b). Specifically, the average degree is 〈*k*〉 ≃ 6 at high odorant concentration (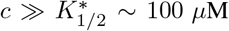 [16]), but is reduced to 〈*k*〉 ≃ 3.6 at 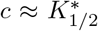. The degree distributions still look qualitatively similar (consistent with the form *P*(*k*) ~ *k*^−*α*^), while the decay becomes steeper at lower concentrations (Fig. S2c). We also show three snapshots of the interaction network at *c* = 0.1M, *c* = 100*μ*M and *c* = 1*μ*M respectively (Fig. S2d-f).

**Figure S1:**
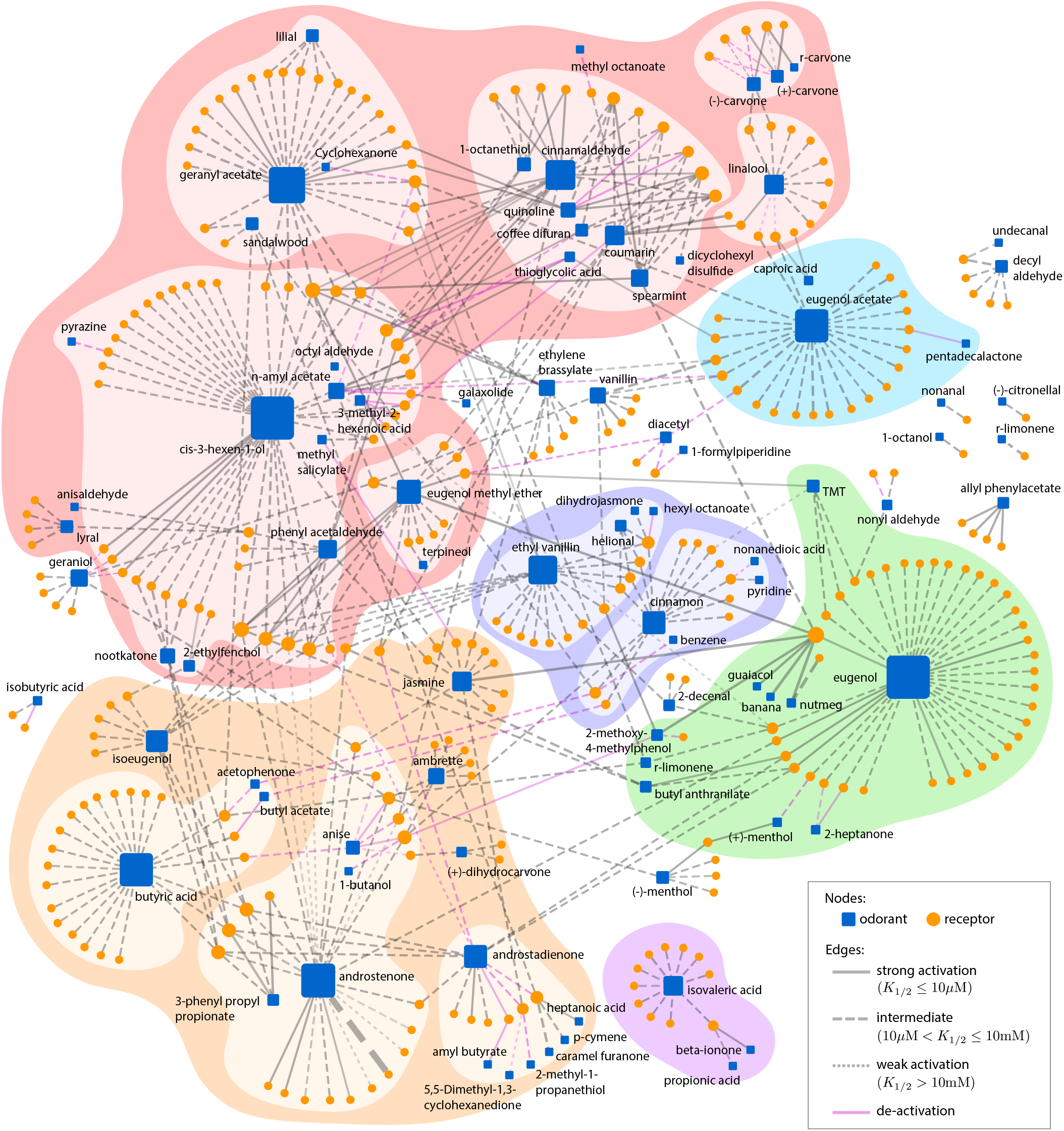
Interaction network with full edge attributes. The edge color indicates the signs of interactions, with gray for activations and magenta for de-activations. The edge line style indicates the range of the characteristic concentration *K*_1/2_: we used solid lines (log_10_ *K*_1/2_ ≤ −5), dashed lines (−5 < log_10_ *K*_1/2_ ≤ −2), and dotted lines (log_10_ *K*_1/2_ > −2), as also shown in the legends in Fig. S1. We note that the distribution of *K*_1/2_ has a peak around the average value 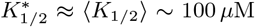 [16]. The edge thickness indicates the net amplitude of the response, *A*, such that a thicker line represents an interaction that elicits a larger response. We also varied the edge transparency to indicate the goodness of fit for the determination of the four parameters, such that a more opaque (less transparent) edge means a higher correlation coefficient, or a better fit. See the accompanying supplementary text for more details. A network file with full information on all nodes and edges, prepared for easy interactive access, is also available online.

**Figure S2:**
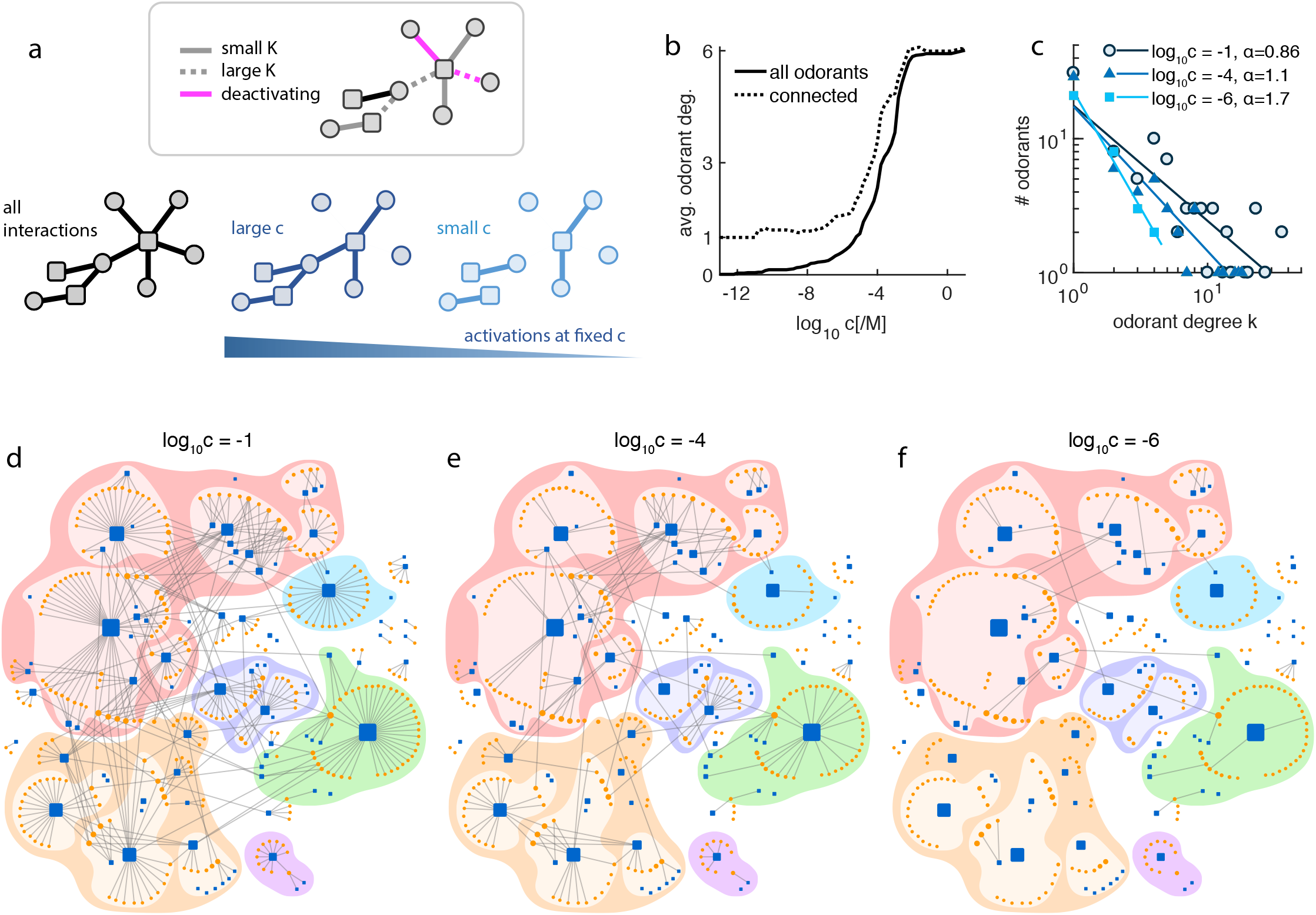
Interaction network at fixed odorant concentrations. **a.** Construction of the binary interaction network at fixed odorant concentration, *c*, assumed uniform for all odorants. **b.** The average degree of odorants at varying *c*. Average was taken either over all the odorants in the network (solid line) or over the odorants connected to at least one receptor at the respective concentration level (dotted line). **c.** Degree distributions obtained by counting activations at three fixed odorant concentrations (*c* =0.1 M, 100 *μ*M, 1 *μ*M), each showing a heavy tail but a different rate of decay. **d-f:** Binary interaction networks at the three fixed odorant concentrations. For easy comparison, node sizes are inherited from the original network; i.e., they reflect the number of all interactions, including the ones that are suppressed at the respective concentrations.

**Figure S3:**
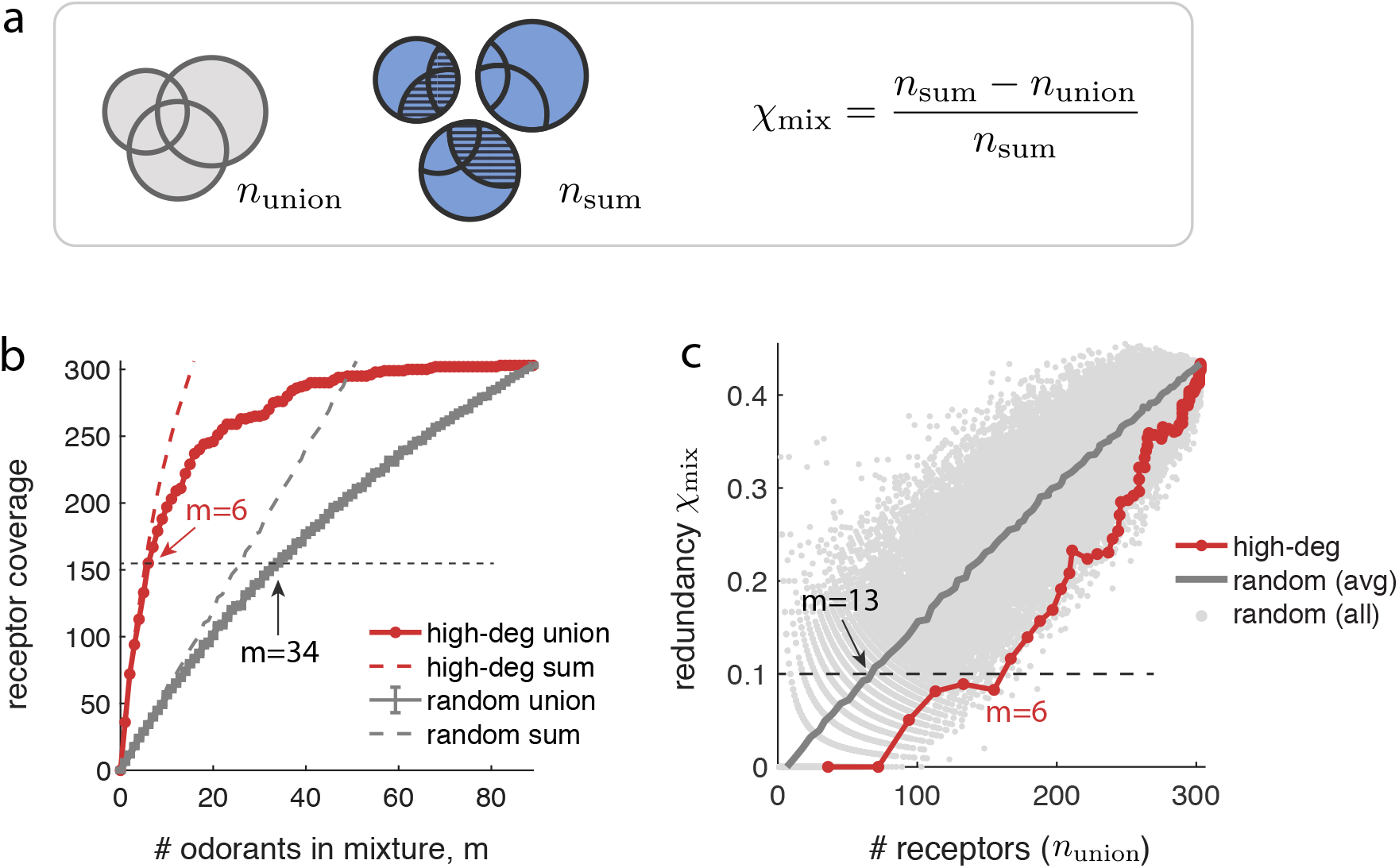
Odorant mixture test. **a.** Illustrations of the net receptor coverage *n*_union_ (gray area, which also equivalent to unstriped blue area), and the sum of individual receptor coverages *n*_sum_ (total blue area), in a mixture of odorants. The receptor code redundancy in a mixture, χ_mix_, is defined by the fraction of the overlap (striped area) with respect to **b.** We plot the receptor coverage by a mixture of odorants, *n*_union_, at varying number of odorants in the mixture, *m*. A mixture contains either the *m* highest-degree odorant in the test dataset (red) or a random selection of *m* odorants from the test dataset (gray); we take 500 independent samples of random mixtures. With the six highest-degree odorants, the receptor coverage is *n*_union_ = 155, which is comparable to a random mixture with *m* ≈ 34 odorants on average. The dashed lines show *n*_sum_ for comparison. **c.** We plot the redundancy in a mixture, χ_mix_, versus the receptor coverage for the corresponding mixture, n_union_. Again, a mixture is selected either from the highest-degree odorants (red) or randomly (gray). The gray line indicates the redundancy calculated from the average counts 〈*n*_union_〉 and 〈*n*_sum_〉 over all 500 random mixtures at each *m*. Gray dots show the results from all 45,000 random mixtures (500 repetitions at each m, and 90 different values of *m*).

**Figure S4:**
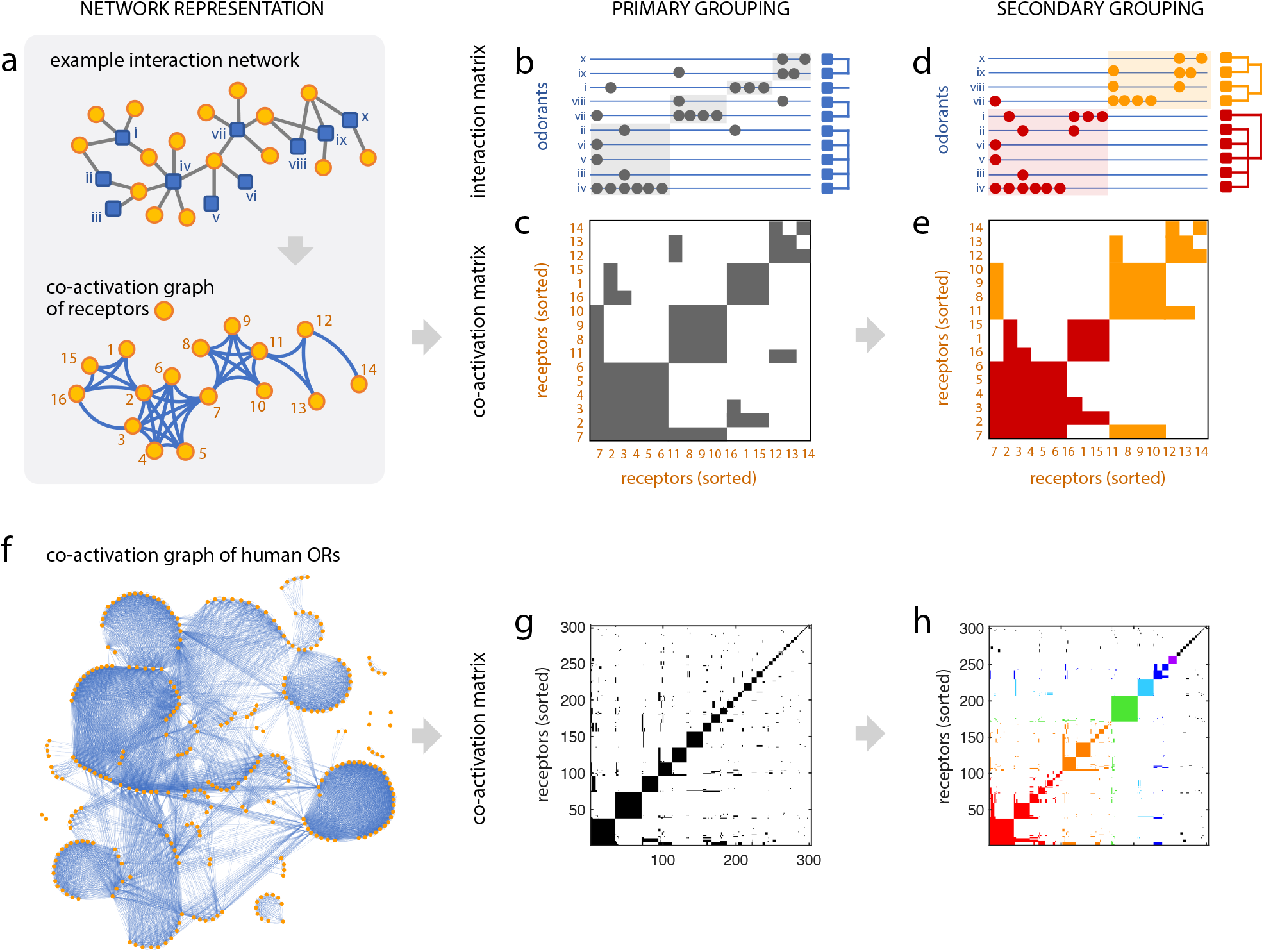
The grouping procedure. **a-e.** Illustration using an example interaction network. **a.** The co-activation graph is a one-mode projection from the bipartite interaction network. Co-activation graph of either node type can be considered, but disjoint hubs (high-degree nodes) in one node type give rise to isolated cliques in the co-activation graph on the other node type. In this example case, almost disjoint odorant hubs (blue squares) are reflected in the almost isolated cliques in the co-activation graph of receptors (orange circles). **b.** Odorants are grouped consequently. **c.** A co-activation matrix, where each row or column is a receptor; receptor cliques appear as square blocks. Receptors are sorted such that cliques are ordered by the size; this is called the primary grouping. **d.** Odorants are again grouped consequently. **e.** Receptors are grouped once again by the redundancy, and re-sorted according to this secondary grouping. See text for details. **f-h.** Analysis on the odorant-receptor interaction dataset [14], binarized by counting all interactions. **f.** The co-activation graph from the real dataset. **g.** Sorted co-activation matrix for the 303 receptors. Diagonal blocks indicate the primary grouping of receptors. each around a shared hub odorant. **h.** Secondary grouping of receptors, based on the receptor code redundancy between the odorant hubs. We colored the 6 largest groups, which already span more than 250 receptors cumulatively. The hub odorants for several large primary groups are labeled.

**Figure S5:**
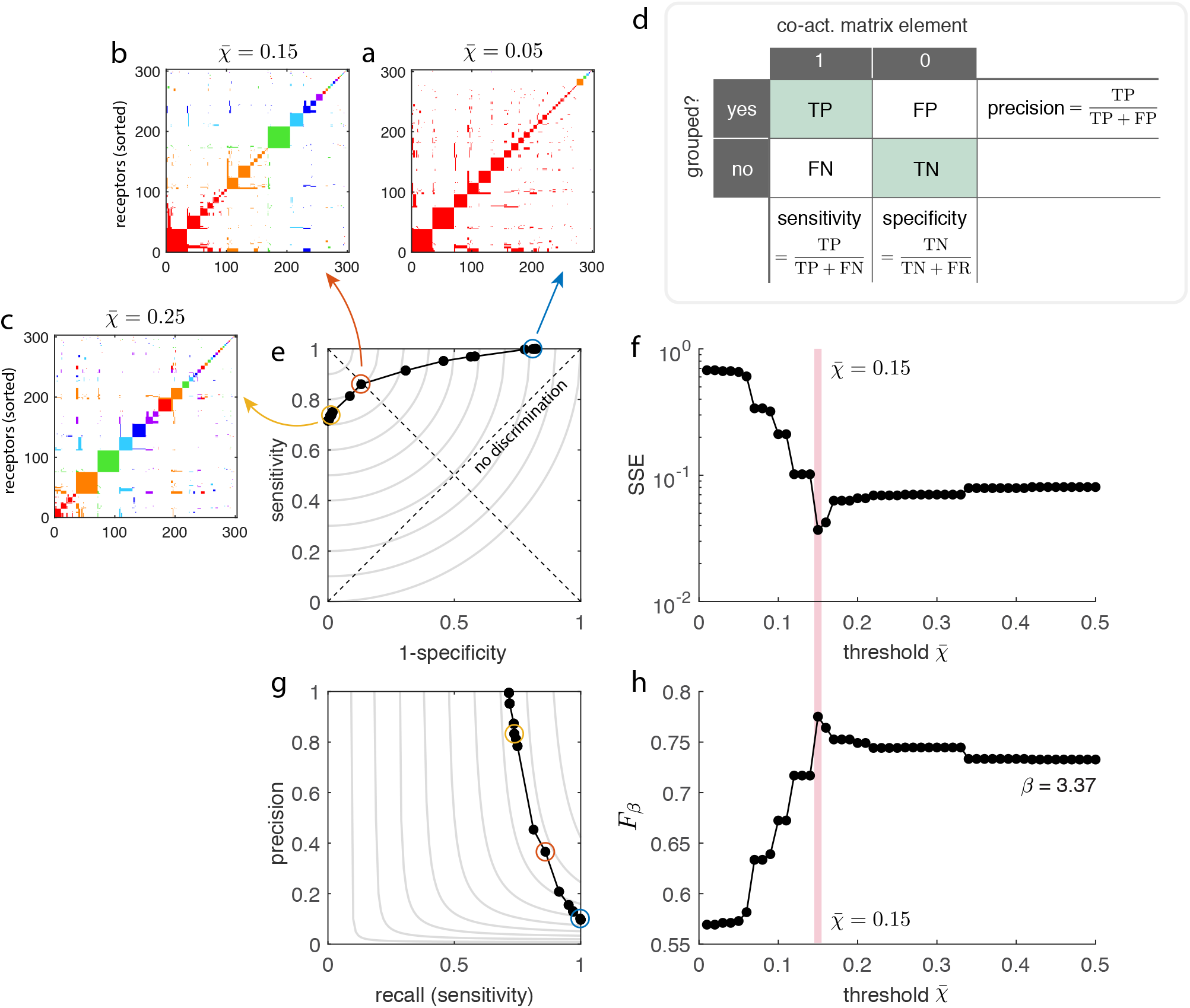
Finding the optimal threshold 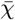 for the secondary grouping of receptors. **a-c.** Secondary grouping results at different thresholds, 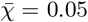, 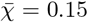 and 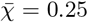 respectively. Each panel shows the co-activation matrix of receptors, sorted and colored by the corresponding secondary groups. **d.** The 2 × 2 contingency table for the binary classifier, and the different definitions of success rates considered in this analysis. **e.** The ROC curve of grouping at varying threshold 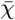. The two diagonals, the line of no discrimination which corresponds to random guesses (positive diagonal) and the line of balance points where sensitivity equals specificity (negative diagonal), are shown in dashed lines. Gray contours indicate the lines of constant error (sum of squared error; SSE). **f.** SSE at varying threshold 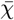. **g.** The precision-recall curve at varying threshold 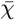. Gray contours indicate the lines of constant *F_β_* score, evaluated at *β* = 3.37 that represents the balance of positives and negatives in data. **h.** *F_β_* at varying threshold 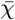.

**Figure S6:**
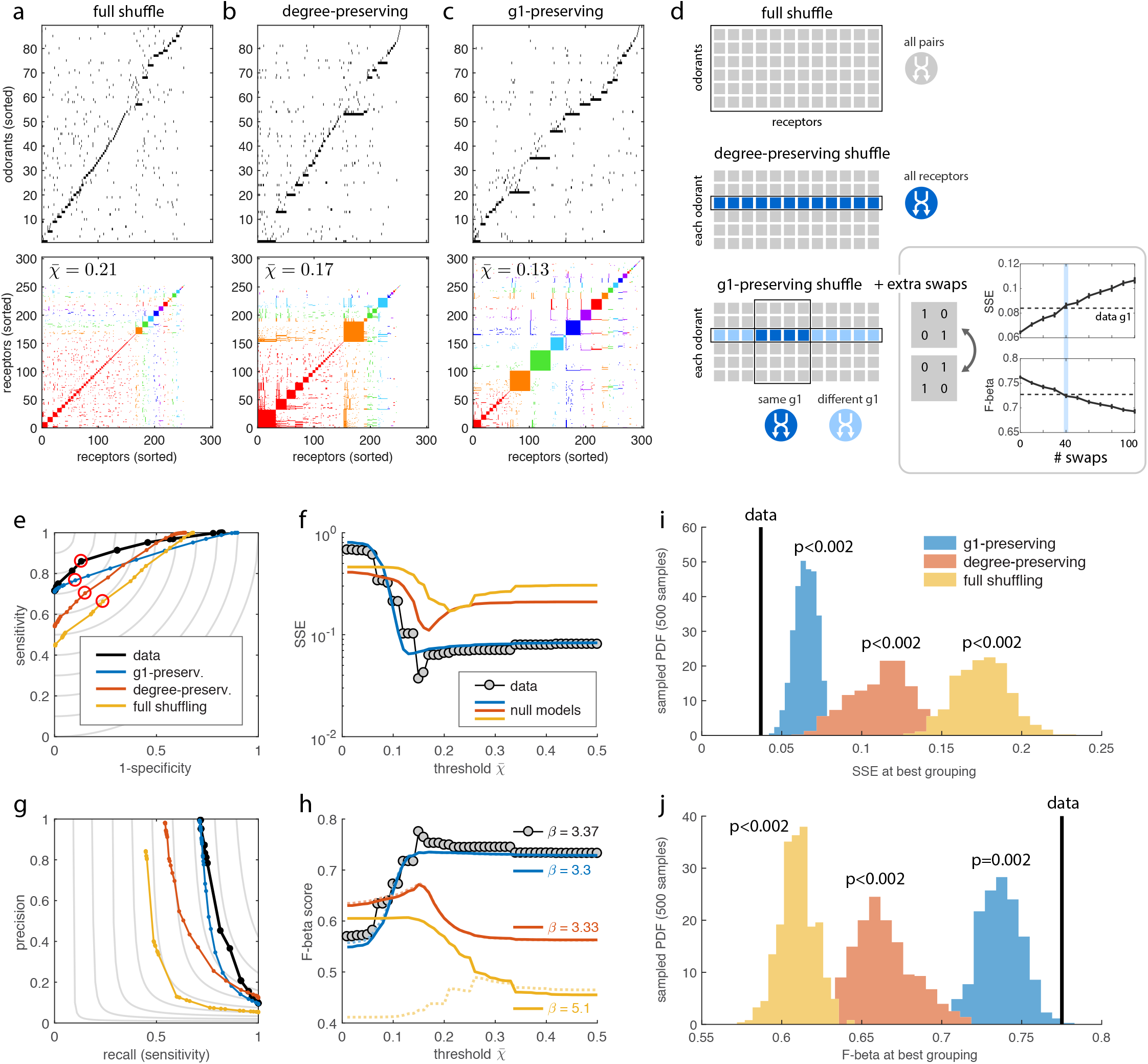
Shuffling test for receptor code grouping. **(a-c)** Examples of null models, obtained by **a.** full shuffling, **b.** degree-preserving shuffling, and **c.** primary group (*g*_1_)-preserving shuffling and extra swaps. Upper panels show the shuffled interaction matrices. Lower panels show the receptor co-activation matrices, sorted and colored by the best grouping, in the sense of prediction error minimization (red circles in **e**). **(d)** Schematic illustrations of the three shuffling rules. See Methods for more details. The goodness of the original primary grouping is restored at 40 extra swaps, for both goodness measures we considered in this analysis (SSE and *F_β_*, also see the next panels). Plots show the averages over 30 repetitions of shuffle-and-swaps, with standard error bars. **(e-f)** Sensitivity-specificity analysis. **e.** The ROC curves for the real data and the three null models, with the lines of constant sum-of-squared error (SSE) shown in gray. **f.** SSE at varying threshold 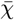. **(g-h)** Precision-recall analysis. **g.** The precision-recall curves, with the constant-*F_β_* lines (evaluated at *β* = 3.37) in gray. **h.** The *F_β_* score at varying threshold 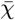. For each null model, the solid line is evaluated with its own value of *β*, whereas the dotted line is evaluated at the *β* = 3.37 determined for the original dataset. **(i-j)** Evaluation of statistical significance by sampling the distributions of the two test statistics, **i.** SSE and **j.** *F_β_* score, under the null models. We sampled 500 independent shuffles under each null model.

**Figure S7:**
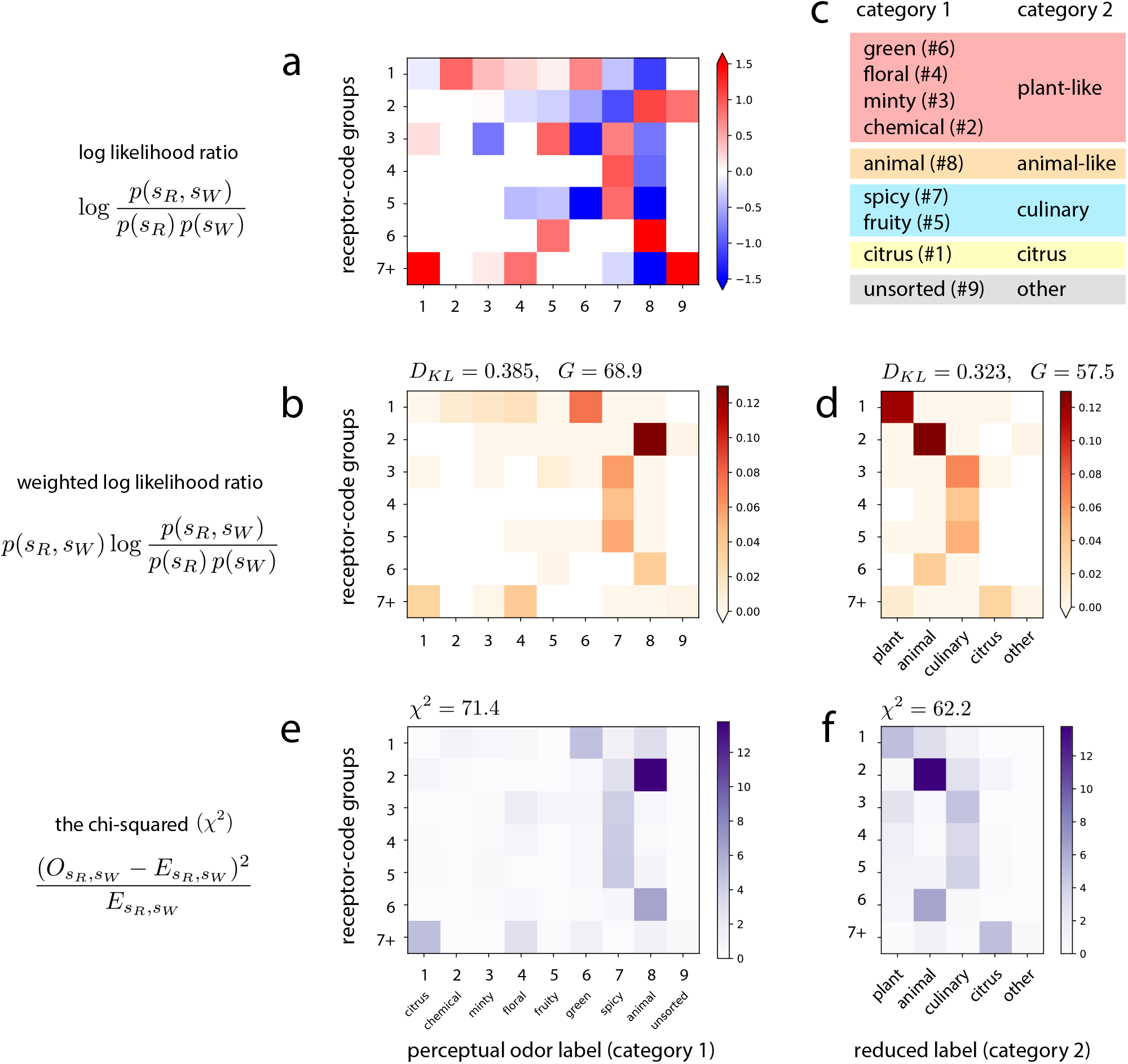
Joint statistics between the receptor-code groups, *s_R_*, and the perceptual odor categories, *s_W_*. **a.** The log likelihood ratios between the full joint distribution (observed) and the uncorrelated null model (expected). **b.** The weighted log likelihood ratios, that sum up to give the Kullback-Leibler divergence *D_KL_* (in this case also equal to the mutual information between *s_R_* and *s_W_*). The corresponding summary statistics are *D_KL_* = 0.385 and *G* = 68.9, which gives *p* = 0.027 with 48 degrees of freedom. **c.** Dimensionality reduction of the perceptual odor categories by merging. **d.** The weighted log-likelihood ratios using the reduced odor categories. The summary statistics are *D_KL_* = 0.323 and *G* = 57.5, which gives *p* = 1.4 × 10^−4^ at 24 degrees of freedom. **e-f.** The local chi-squared values. **e.** Using all eight perceptual odor categories, we get χ^2^ = 71.4 and *p* = 0.016 (48 degrees of freedom). **f.** Using the reduced categories, we get χ^2^ = 62.2 and *p* = 3.0 × 10^−5^ (24 degrees of freedom).

**Table S1:**
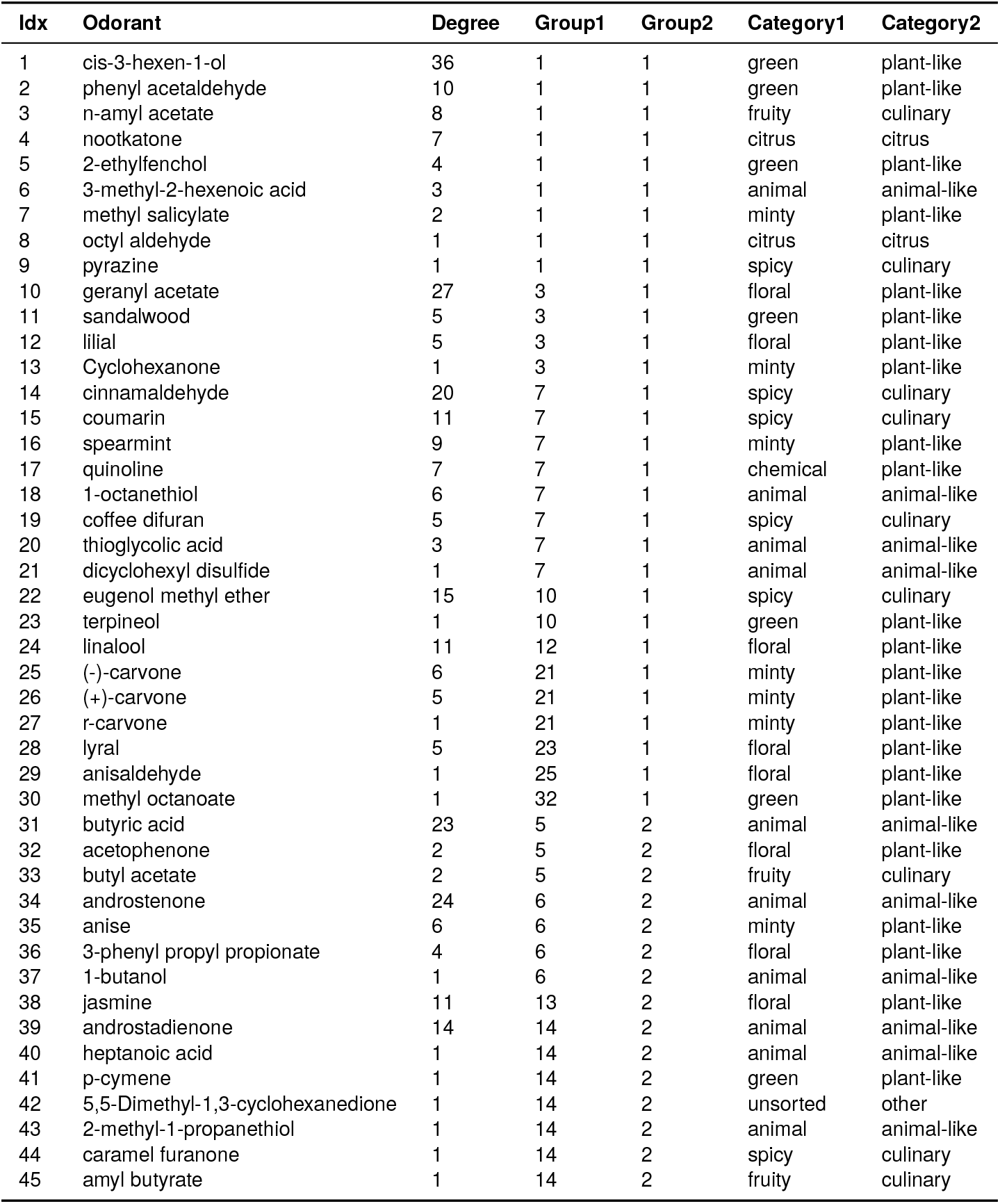
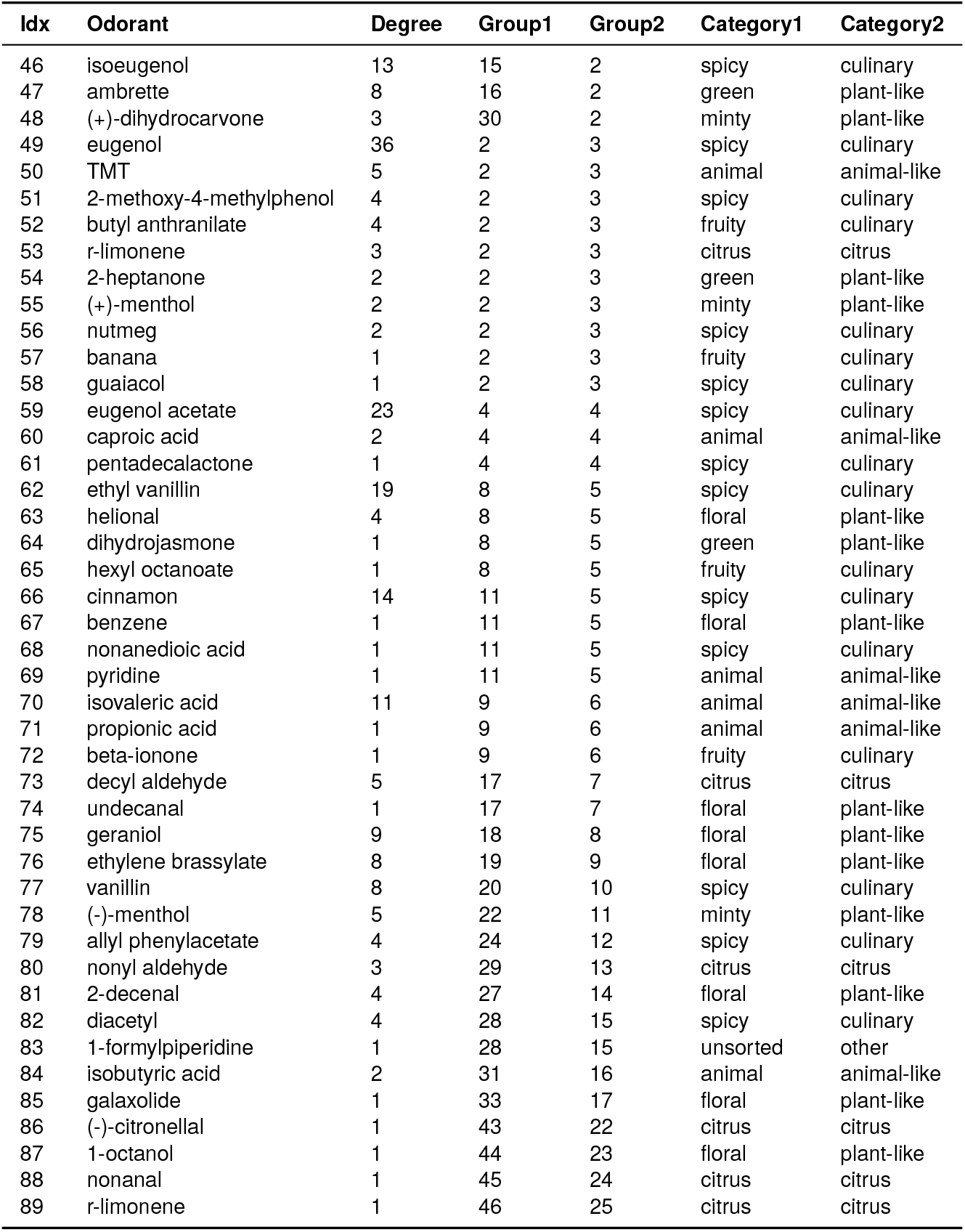
List of odorants with receptor-code group and perceptual odor category labels. List of odorants with receptor-code group and perceptual odor category labels. The 89 odorants are listed according to the groups identified in our work. The order of odorants is the same as in the row order in Fig. 4 in the main text. The columns “Group1” and “Group2” show the primary odorant group rank and the secondary odorant group rank, respectively, ranked by the size of the corresponding receptor groups. The columns “Category1” shows the perceptual odor category as adopted from [21]. “Category2” shows the reduced odor label, obtained by merging Category1.

**Table S2:**
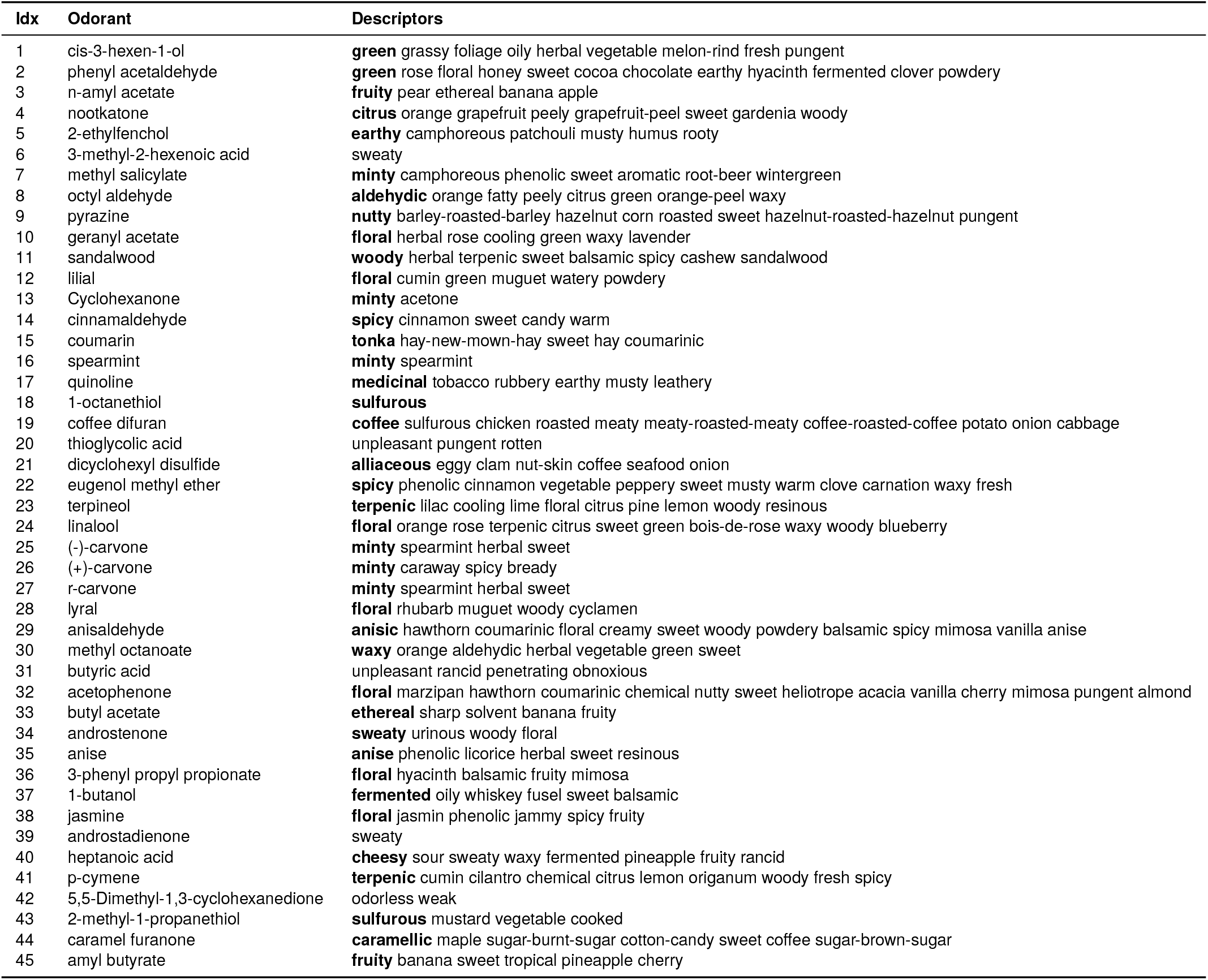
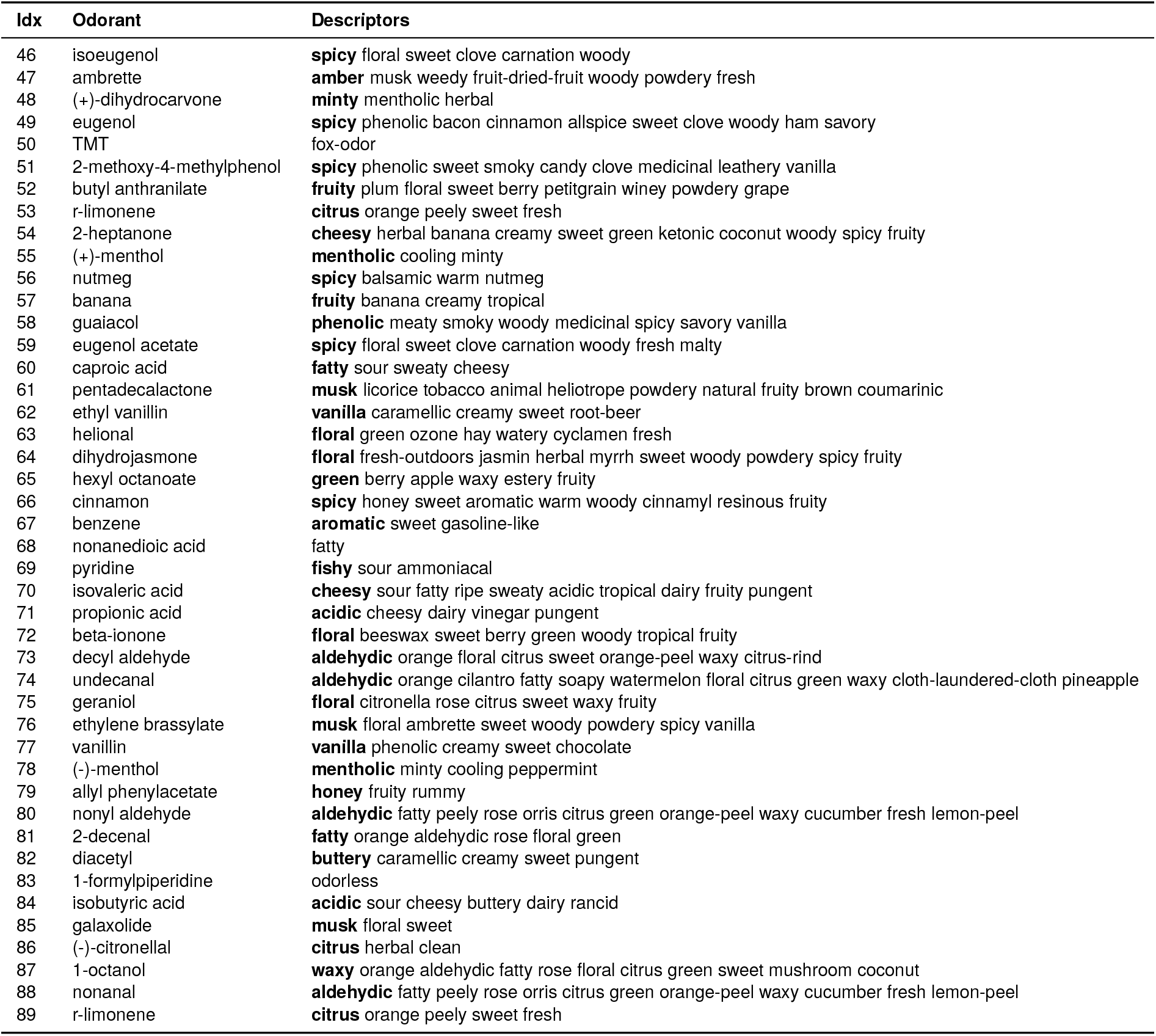
List of odorants with perceptual odor descriptors. For each odorant, all descriptors were obtained from a single source. If they were obtained from the Good Scents Company Information System [56], and if an odor type was specified, the odor type descriptor is boldfaced in the list. Descriptor sources are separately provided in Table S5.

**Table S3:** Reference descriptors for the perceptual odor categories. Category numbers and the list of reference words are copied directly from Figure 7 in [21]. We name each category for convenience (column Nickname).

| Category | Nickname | Reference |
| --- | --- | --- |
| 1 | citrus | fruity/citrus, lemon, grapefruit, orange |
| 2 | chemical | nail polish remover, mothballs, alcoholic, etherish, anaesthetic, cleaning fluid, gasoline, solvent, turpentine (pine oil), leather, tar, creosote, disinfectant, carbolic, medicinal, chemical, ammonia, new rubber, kerosene, paint, varnish, metallic |
| 3 | minty | tea leaves, caraway, minty, peppermint, camphor, eucaliptus, anise (licorise), cheesy, cool, cooling |
| 4 | floral | floral, rose, violets, lavender, cologne, musk, perfumery, fragrant, aromatic, soapy, incense, light |
| 5 | fruity | fruity/other than citrus, pineapple, grape juice, strawberry, pear, cantaloupe, honeydew melon, peach (fruit), banana, cherry (berry), sweet, raisins |
| 6 | green | laurel leaves, black pepper, green pepper, dill, oak wood, cognac, woody, resinous, cedarwood, geranium leaves, celery, fresh green vegetables, crushed weeds, crushed grass, herbal, green, cut grass, raw cucumber, cardboard, rope, wet paper, musty, earthy, moldy, raw potato, mushroom, beany, bark, birth bark, cork, dry, powdery, chalky |
| 7 | spicy | honey, almond, nutty (walnut etc), spicy, clove, cinnamon, chocolate, vanilla, maple syrup, caramel, malty, molasses, coconut, hay, bakery (fresh bread), peanut butter, burnt, smoky, fresh tobacco smoke, coffee, stale tobacco smoke, burnt paper, burnt milk, burnt candle, oily, fatty, buttery, fresh butter, popcorn, fried chicken, warm |
| 8 | animal | apple (fruit), seasoning (for meat), grainy (as grain), yeasty, sour milk, fermented (rotten), fruit, beery, wet wool, wet dog, dirty linen, stale, mouse, eggy (fresh eggs), burnt rubber, bitter, sharp, pungent, acid, sour, vinegar, sauerkraut, urine, cat urine, fishy, kippery (smoked fish), seminal, sperm, sooty, meaty (cooked, good), soupy, cooked vegetables, rancid, sweaty, household gas, sulfidic, garlic, onion, blood, raw meat, animal, sewer, putrid, foul, decayed, fecal (like manure), cadaverous (dead animal), sickening, heavy |

**Table S4:**
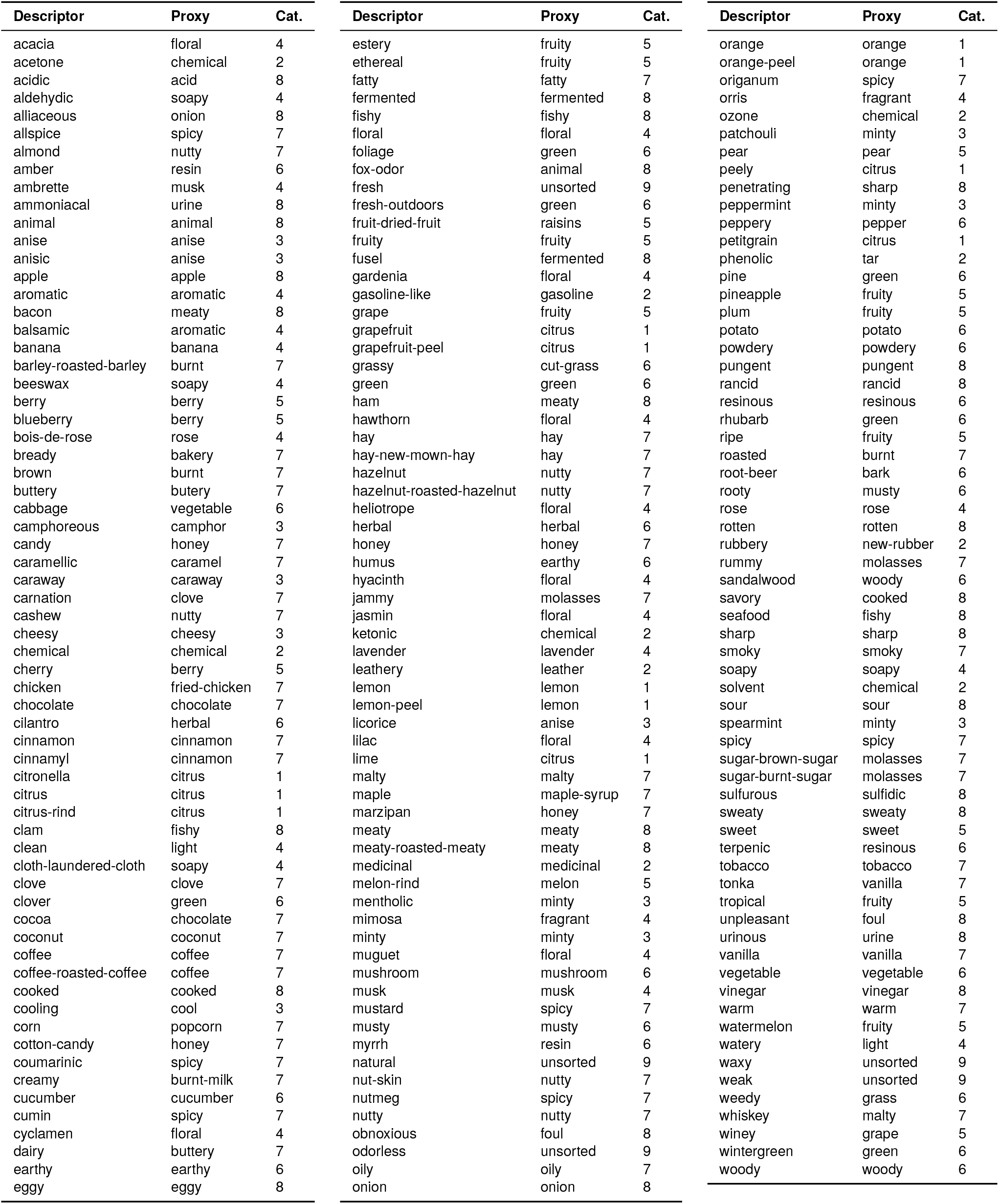
Labeling odor descriptors. For each of the 184 test descriptors, we chose a best approximation (proxy) from the list of reference words for the perceptual odor categories (Table S3). The category label is determined by the proxy.

**Table S5:**
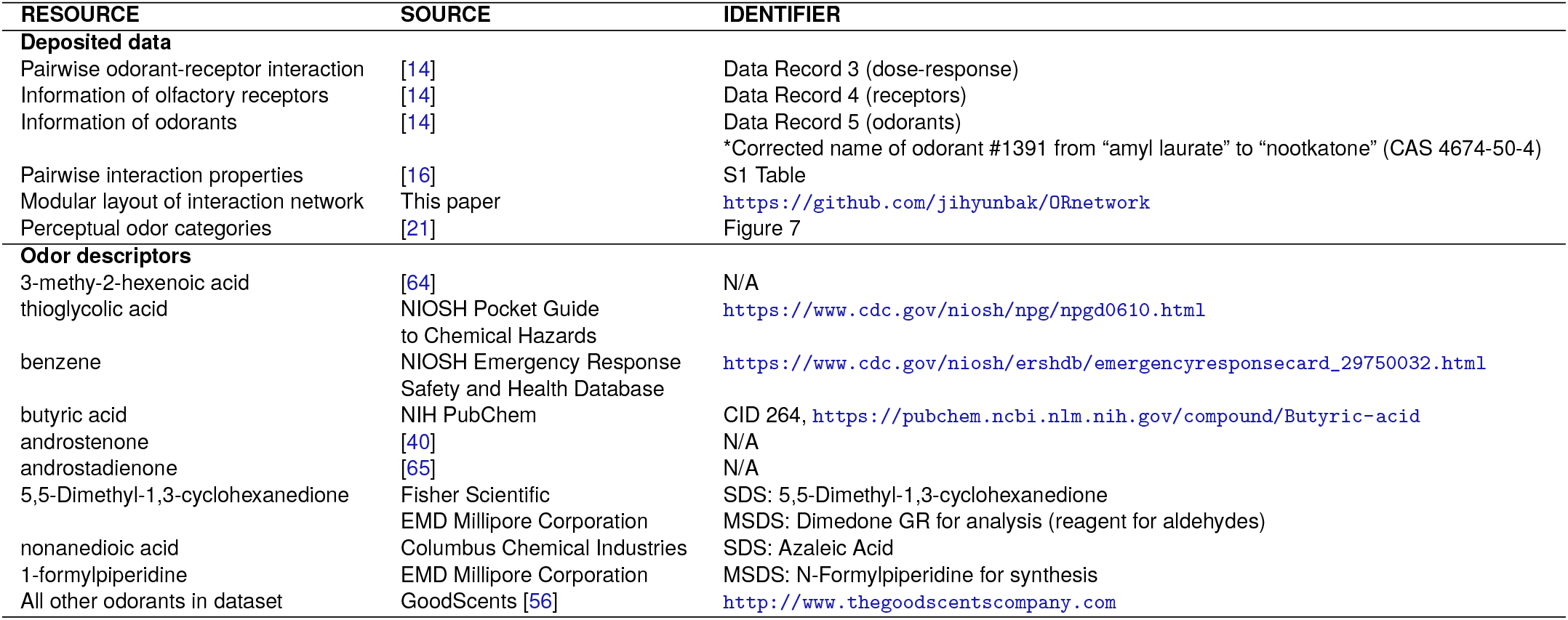
List of data sources. SDS: Safety Data Sheet; MSDS: Material Safety Data Sheet.

